# Matching cell lines with cancer type and subtype of origin via mutational, epigenomic and transcriptomic patterns

**DOI:** 10.1101/809400

**Authors:** Marina Salvadores, Francisco Fuster-Tormo, Fran Supek

**Affiliations:** Institute for Research in Biomedicine (IRB Barcelona), The Barcelona Institute of Science and Technology, Barcelona, Spain; MDS Research Group, Institut de Recerca Contra la Leucèmia Josep Carreras, Institut Català d’Oncologia-Hospital Germans Trias i Pujol, Universitat Autònoma de Barcelona, Badalona, Spain; Institució Catalana de Recerca i Estudis Avançats (ICREA), Barcelona, Spain

**Keywords:** cancer cell lines, human tumors, mRNA level, DNA methylation, mutational signatures, cancer type, drug screen, genetic screen

## Abstract

Cell lines are commonly used as cancer models. Because the tissue and/or cell type of origin provide important context for understanding mechanisms of cancer, we systematically examined whether cell lines exhibit features matching the cancer type that supposedly originated them. To this end, we aligned the mRNA expression and DNA methylation data between ∼9,000 solid tumors and ∼600 cell lines to remove the global differences stemming from growth in cell culture. Next, we created classification models for cancer type and subtype using tumor data, and applied them to cell line data. Overall, the transcriptomic and epigenomic classifiers consistently identified 35 cell lines which better matched a different tissue or cell type than the one the cell line was originally annotated with; we recommend caution in using these cell lines in experimental work. Six cell lines were identified as originating from the skin, of which five were further corroborated by the presence of a UV-like mutational signature in their genome, strongly suggesting mislabelling. Overall, genomic evidence additionally supports that 22 (3.6% of all considered) cell lines may be mislabelled because we predict they originate from a different tissue/cell type. Finally, we cataloged 366 cell lines in which both transcriptomic and epigenomic profiles strongly resemble the tumor type of origin, designating them as ‘golden set’ cell lines. We suggest these cell lines are better suited for experimental work that depends on tissue identity and propose tentative assignments to cancer subtypes. Finally, we show that accounting for the uncertain tissue-of-origin labels can change the interpretation of drug sensitivity and CRISPR genetic screening data. In particular, in brain, lung and pancreatic cancer cell lines, many novel determinants of drug sensitivity or resistance emerged by focussing on the cell lines that are best matched to the cancer type of interest.

## Introduction

Cell lines are an important research tool, often used in place of primary cells and intact organisms to study biological processes. Cell lines are used for various applications such as testing drug metabolism and cytotoxicity, study of gene function, generation of artificial tissues and synthesis of biological compounds (1). In cancer research, cell lines derived from tumors are commonly used as models because they are presumed to carry the genomic and epigenomic alterations that arise in tumors (2). To understand the response of tumors to therapy, many studies have linked genetic and/or epigenetic alterations with drug response across cell line panels, generating datasets such as the Genomics of Drug Sensitivity in Cancer (GDSC) (3), the Cancer Cell Line Encyclopedia (CCLE) (4), the Cancer Therapeutics Response Portal (5) and others. These efforts have advanced our understanding of tumor biology by generating a massive resource of genomic, transcriptomic, epigenomic and drug response data for hundreds of cell lines (2).

As a model for cancer, cell lines are cost effective, convenient and amenable to high-throughput screening (1,2). However, a major question associated with the use of cell lines is whether they are representative of the cancer they are meant to model, which may be complicated by issues of misidentification (1,2,6).

Misidentified cell lines may lead to inconsistent conclusions across studies using the affected cell lines. For instance, the cell lines referred to as HEp-2 and INT 407 in the literature are in fact commonly cross-contaminated with HeLa (cervical cancer) cells, rather than being laryngeal cancer and normal intestinal epithelium cells, respectively (7,8). Because of this, demonstrating cell line identity via genetic markers is now a routine quality-control step. Current resources based on large-scale cancer cell panels are therefore largely unaffected by this issue (4).

However, even if the identity of the cell line is correct, its properties may not match the cancer type it is meant to model. One way in which this may happen is that tumors thought to originate in a certain tissue might in fact be metastatic lesions originating from a distal site (9). Thus, cell lines derived from such tumors would have a different tissue/cell type identity than that assigned at isolation, constituting a case of mislabeling. It is conceivable that also in the case of primary tumors, ambiguous histological or anatomical features may cause the cancer type or subtype to be misdiagnosed for that tumor and therefore also for a cell line derived from it. Conceivably, the process of establishing the culture might select for a rare cell type that is not representative of the tumor isolate on the whole, meaning that the cell line would again effectively be mislabeled with a different cell type (10). In addition to the initial changes upon adaptation to culture, cell lines evolve over time due to selection and due to genetic drift, potentially diverging from the characteristics of the originating tissue (1).

Tissue/cell type is a key determinant of response of cultured cells to a variety of experimental conditions, including drug exposure and genetic perturbation (11,12). Therefore, having accurate information on the tissue and cell type identity of a tumor cell line is important for interpreting the experimental results obtained using these cell lines.

Recent work has examined cell line panels of individual cancer types, showing certain discrepancies between the features of cell lines and corresponding tumor (sub)types. A gene expression analysis of lung tumors and cell lines (10) suggested that some lung adenocarcinoma cell lines did not resemble adenocarcinoma tumors but instead clustered with other lung tumor subtypes (small-cell and squamous cell). A study of high-grade serous ovarian cancer (HGSOC) cell lines using gene expression, driver gene mutations and copy number alteration (CNA) data reported that two frequently used cell lines showed poor genetic similarity to profiles of this ovarian cancer subtype (13). A study of a panel of renal cancer cell lines compared their CNA to kidney tumors, finding that some cell lines used as models of the clear-cell carcinoma more closely resemble papillary renal cancer (14). These examples highlight the need to systematically identify the cell lines whose genotype and/or molecular phenotypes do not resemble the characteristics of the matched human tumor type. A major challenge in the use of human tumor data to classify cell lines are the widespread global changes in gene regulation between cell lines and tumors that arise in cell culture conditions.

In this study, we performed a global analysis that aligned mRNA expression and DNA methylation data between ∼600 cancer cell lines and ∼9,000 tumors from 22 different cancer types, adjusting for global differences in transcriptomes and epigenomes. Classifiers trained on human tumor mRNA and DNA methylation profiles were used to systematically identify those cell lines whose genomic and epigenomic profiles are highly consistent with human tumors of their declared cancer type of origin. Conversely, we used the same classifiers to identify those cell lines that might be mislabeled with respect to cancer type or that might have diverged from their original tissue and/or cell type identity. Our data suggests that tens of cell lines might be epigenetically and/or genetically not consistent with their stated tissue or cell type of origin, which is an important consideration for experiments that use these cell lines. We demonstrate this by reanalyzing associations between drug sensitivity and genetic variation in a large panel of cell lines. After explicitly accounting for putative cases of cell lines with mislabeled tissue identity, many novel associations of genes with drug sensitivity or resistance were revealed.

## Results

### 1. Identification of tissue/cell type-of-origin for cell lines by a joint analysis with tumors

During adaptation to cell culture, certain changes in the cell lines’ physiology are inevitable, yet ideally the cell lines should retain sufficient features of the tumor to be useful as experimental models of tumor biology. Here, we systematically examined the global features of the transcriptome and epigenome that reflect the tissue-of-origin of a tumor cell line. The tissue that originated a tumor is well known to be a major determinant of drug responses -- including drugs targeted to certain genetic mutations -- both *in vitro* (11,12) *and* also *in vivo* (15,16). Tissue of origin is an important factor in shaping the networks of genetic interactions in cancer (17) and also determines the phenotypes resulting from genetic perturbation (18). Therefore ascertaining the tissue/cell type identity of cell lines is relevant for interpreting results of various experiments.

During the process of adaptation to cell culture, the cells undergo global changes in gene regulation that affect many genes (19,20). In particular, the global patterns in transcriptomes and epigenomes for cultured cells bear many similarities to other cultured cells, irrespective of the originating tissue. Thus, there are commonalities in how culture affects gene regulation: for example, proliferation genes in cultured cells have distinct DNA methylation and gene expression patterns, when compared to tumor and normal tissues (19,20). These global alterations in gene expression and DNA methylation mean that is not straightforward to directly compare cell line transcriptomes/epigenomes with data obtained from actual tumors. Therefore, the cell culture-induced shifts need to be carefully adjusted for in order to be able to track down tissue identity of cell lines. To this end, we introduce a computational framework -- HyperTracker -- which can unify transcriptome, epigenome and mutational data across tumors and cell lines, and provide robust predictions of tissue/cell type and subtype identity.

In particular, we collected gene expression (RNA-Seq) and DNA methylation data (microarrays) for 9,681 and 9,039 human tumors, respectively (TCGA), and additionally for 614 cell lines (CL) of various solid cancer types. For gene expression data (GE), we examined transcript-per-million (TPM) normalized counts for the 12,419 genes where RNA-Seq data was available for both cell lines and tumors. For DNA methylation data (MET), we examined beta-values for 10,141 probes from methylation arrays, after selecting a single probe per gene promoter with the highest variance across the dataset. To align human tumor and cell line data, we quantile-normalized the data and applied ComBat, a batch effect correction method (21), which is highly performant compared to other related methods (22). In brief, ComBat estimates parameters for location and scale adjustment of each batch (TCGA and CL in our case) for each gene. Then, it removes the variability which is particular to the CL but not present in TCGA, while retaining the intra-dataset variability of the tumors, which should presumably be evident in both the tumor and in the cell line datasets.

A principal component analysis in the data (pre- and post-adjustment) suggests that there were indeed strong global differences between TCGA and CL, and that they are largely removed by our approach (Fig 1a; Fig S1ab). To quantify this, we trained a classification model that predicts the CL *versus* TCGA origin of the data points based on GE and MET (Fig 1b). The model is able to distinguish CL *versus* TCGA perfectly when using the pre-adjustment datasets (AUC=1), while the post-adjustment datasets (AUC(GE) = 0.44; AUC(MET) = 0.42) do not perform better than random (0.5; Fig 1b), suggesting the cell-type specific signal has been largely removed. Finally, we tested the optimal number of features (genes/probes) using tumor classifiers and calculating the accuracy in the cell line data (Fig S1c); we selected 5,000 features with the highest standard deviation for later analyses.

**Fig. 1.**
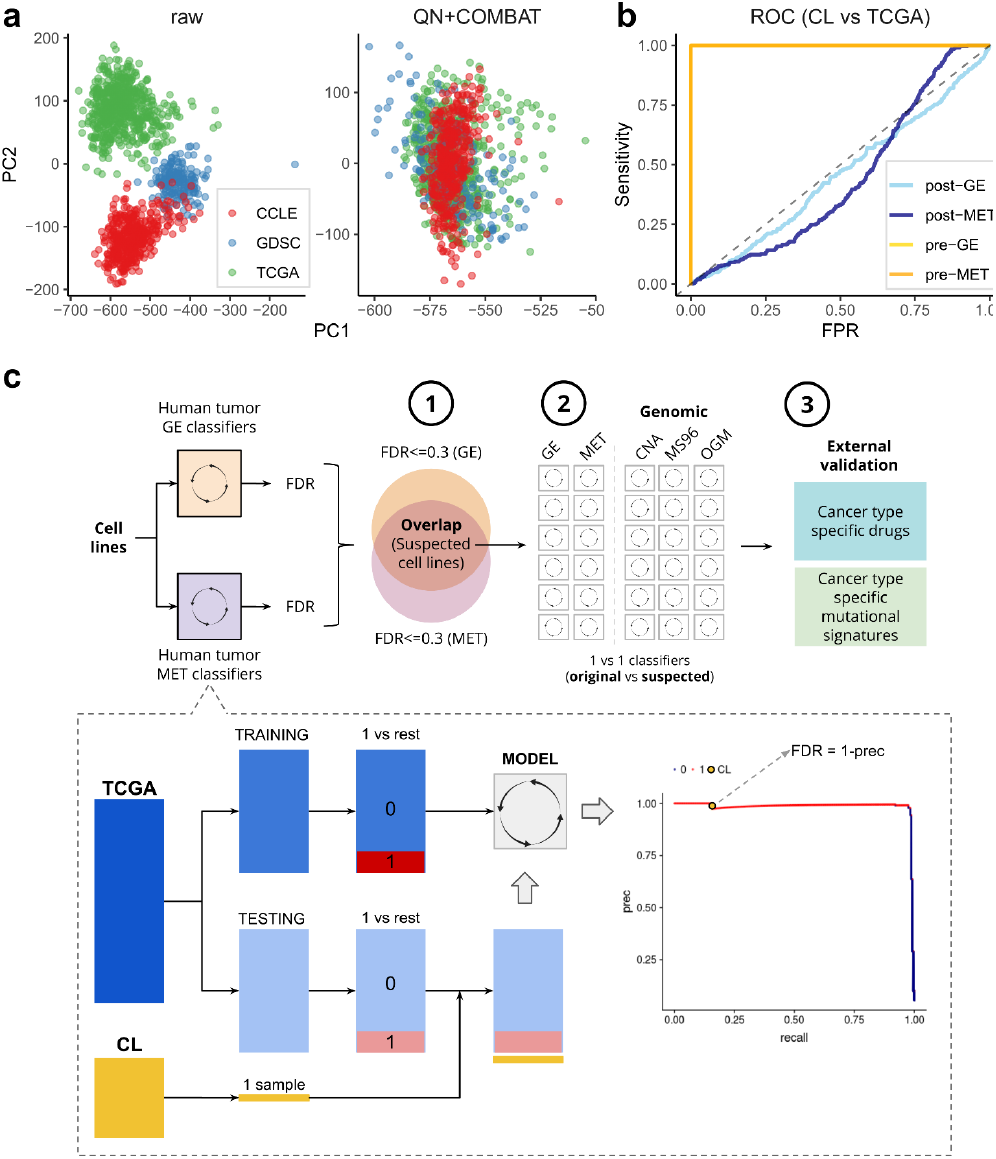
Data alignment and methodology for classification. **(a)** Principal component (PC) 1 and PC2 of a PC analysis, in the gene expression (GE) data pre-adjustment for batch effects (raw) and post-adjustment (quantile normalization+COMBAT) (see Fig S1 for DNA methylation data (MET)). Colors represent the dataset sources (GDSC and CCLE are two sources for the cell lines, and TCGA is the source for the tumors). **(b)** ROC curves for classifying TCGA versus cell lines in the data pre-adjustment (orange) and post-adjustment (blue) for GE and MET. **(c)** An overview of the HyperTracker methodology applied in the manuscript. First, we systematically identified possible mislabeled cell lines using GE and MET data, independantly. Second, we used various types genomic data to corroborate the hits. Third, we further validate the cell lines suspected to originate from skin using independent data.

Once the data was aligned, we set out to determine which cell lines have tissue identity not matching the declared tissue-of-origin (henceforth: ‘suspect set’), and conversely, which cell lines have largely retained their tissue identity (henceforth: ‘golden set’), by comparing against a large set of tumors from 17 tissues in the TCGA (Fig 1c). Using TCGA data, we derived one-*versus*-rest classification models (using Ridge regression), separately for the GE and the MET data. Some pairs of cancer types were considered jointly in this analysis, based on their overall similarity, for example stomach adenocarcinoma (TCGA code: STAD) and esophageal adenocarcinoma (subset of samples from TCGA code: ESAD); see Methods for a full list. Our study focuses on solid cancer types and does not examine blood cancers. In a crossvalidation test, TCGA models had very high AUPRC scores: 0.98 and 0.97 for GE and MET respectively (average across cancer types). This means that transcript level data and DNA methylation data are largely sufficient to accurately distinguish those cancer types.

Next, we obtained predictions of cancer type identity for each cell line. For every cancer type, we split TCGA data randomly into training and testing sets, and we used the calculated precision-recall curve of the testing data to obtain the False Discovery Rate (FDR) score for every cell line (details in Methods; all FDR values are listed in Table S1). The smaller the FDR, the more likely the cell line is to belong to that particular cancer type. As expected, most of the cancer type labels of the cell lines match the declared tissue of origin of that cell line -- they tend to cluster at low FDR values for the cognate cancer type (red dots in Fig 2a, Fig S2). However, among these many correctly classified cell lines (red dots), there are some with similarly low FDR scores, but which were originally annotated as belonging to another cancer type (Fig 2a; blue dots with label shown). A clustering analysis of the GE and MET values for the genes that had the highest weight in the classification models (Fig 2b, Fig S3) showed that in most cases, the samples clearly cluster by cancer type, but not by CL *versus* TCGA label. Moreover, we observed that the suspected cell lines (cell lines with highly confident FDR scores to a different cancer type) tend to cluster with the newly-assigned cancer type by the classifier, rather than with the original one (Fig 2b).

**Fig. 2.**
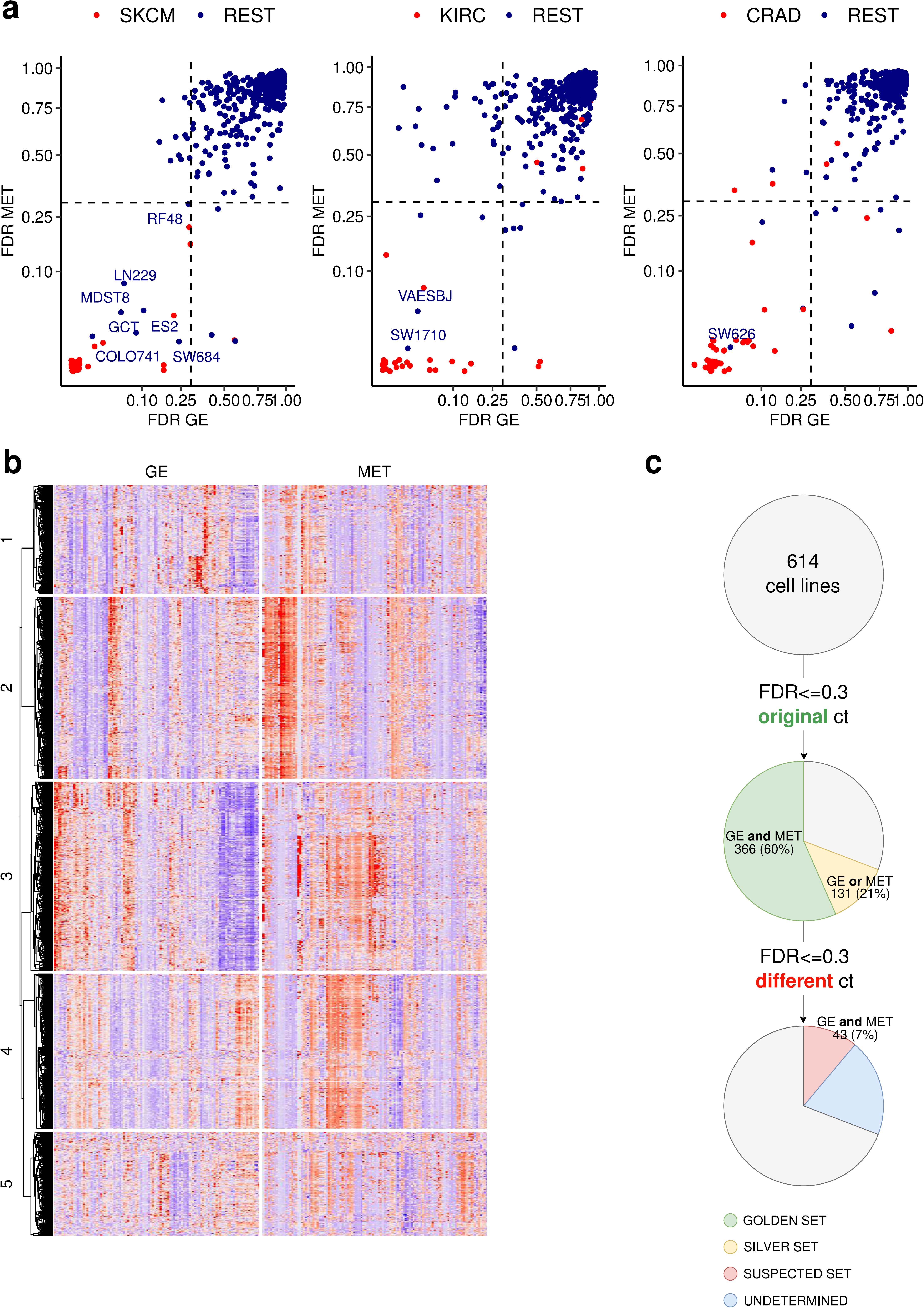
Detection of cell lines mislabelled to a different cancer type. **(a)** False Discovery Rate (FDR) scores for 614 cell lines were calculated in MET and GE cancer type classifiers (one-versus-rest). The lower the FDR, the higher the confidence that the sample belongs to that particular cancer type (here, to SKCM, KIRC and CRAD from left to right, see Fig S2 for the other cancer types). The cell lines that were originally annotated as the cancer type that is being tested are shown in red, the rest in blue. **(b)** Heatmap for the 25 genes (GE) and CpG probes (MET) whose Ridge regression coefficients had the highest absolute values for SKCM (skin cancer) versus rest classifiers. The suspected skin cell lines are labeled in the right side of the heatmap. The cancer types shown are the suspected one (SKCM in this case) and additionally the originally declared cancer types of the suspected cell lines (here, ESTAD, SARC, CRAD and GYNE). See Fig S3 for the Heatmaps for the rest of the suspected cell lines. **(c)** Overview of the results from the systematic mislabelling testing of all cell lines. Cell lines with an FDR<=0.3 to its original cancer type in (i) GE and MET are assigned to the ‘golden set’ group and (ii) either GE or MET are assigned to the ‘silver set’. If however the FDR<=0.3 to a different cancer type in GE and in MET, the cell line is assigned to the suspected set.

In further analyses, we designated as the ‘golden set’ those cell lines that have FDR <= 0.3 for both GE and, independently, for MET in their originally declared cancer type (*n*=366 out of 614 examined cell lines, 60%). For these cell lines, two independent types of evidence -- transcriptomes and epigenomes -- support that they match their expected cancer type well, suggesting these cell lines would be preferred as experimental models. Further, we designated as the ‘silver set’ those cell those cell lines that have FDR <= 0.3 for only one classifier (either GE or MET but not both) (*n*=131 out of 614 examined cell lines, 21%). From the remaining 117 cell lines, we selected as ‘suspect set’ those CL which exhibit an FDR <= 30% for both GE and for MET, but in a different cancer type than declared for that cell line (n=43 out of 614, 7% of analyzed cell lines) (Fig 1c). This set of cell lines may consist either of mislabeled cell lines, where the cancer type of origin is different than it was thought, or of heavily diverged cell lines, where the genomic and/or epigenomic alterations accumulating during cell culture have overridden the original cancer type identity. Of note, cell line cross-contamination issues (23) cannot underlie the trends we observe, because the repositories that provided GE and MET data have used genetic markers to ascertain the identity of the cell lines (4). The fact that two classifiers based on independent data types -- one genomic and one epigenomic -- reached the same predictions adds confidence that these are *bona fide* cases of mistaken tissue/cell-type identity.

### 2. Validation of individual examples of suspected mislabeled cell lines using genomic classifiers

We detected 43 cell lines that bear transcriptomic and also epigenomic features of a different cancer type than the one they were originally annotated to. We next turned to support individual examples of cell lines with reassigned tissue identity by analyzing independent data. In particular, we used genomic sequence-based classifiers, which are able to predict the tissue of origin based on mutation patterns (24,25). As in our recent work (24), we used the trinucleotide mutation spectra and the oncogenic mutations. In this validation setting, we applied such genomic classifiers to a problem of ‘one-versus-one’ classification, where we contrasted the originally assigned cancer type *versus* the newly-proposed cancer type for each reassignment. We found that such one-versus-one classifiers based on genomic data had satisfactory accuracy with our whole-exome sequencing data sets (Fig S4; our past work (24) suggests whole genome sequences are more powerful). Finally, we included an additional classifier based on copy number alteration (CNA) profiles, which were also shown to yield accurate predictive models of tissue specificity (24,25).

For the 43 examples of suspected cell lines tissue identity, we first derived one-*versus*-one classification models separately for GE and MET. If a cell line is truly mislabelled when testing the original *versus* the suspected cancer type, we should observe the same reassignment of the cell line to be robustly observed across multiple runs of the classification algorithm, which use different random initializations. Out of 20 iterations of the algorithm, a score of 20 indicates that the cell line is consistently predicted as the suspected cancer type, and a score of 0 means that the cell line is consistently assigned to the original cancer type. We randomized the labels to obtain a background model of expected values (Fig 3b; Fig S5a). From the 43 suspected cell lines, 35 are consistently reassigned to the other tissue (score>10), irrespective of the variability in the predictive models introduced by resampling the data (Fig 3a; Fig S5b). Next, we calculated the same score for the genomic classifiers (based on mutations and CNA, as described above) on these 35 suspected cell lines (Fig 3a).

**Fig. 3.**
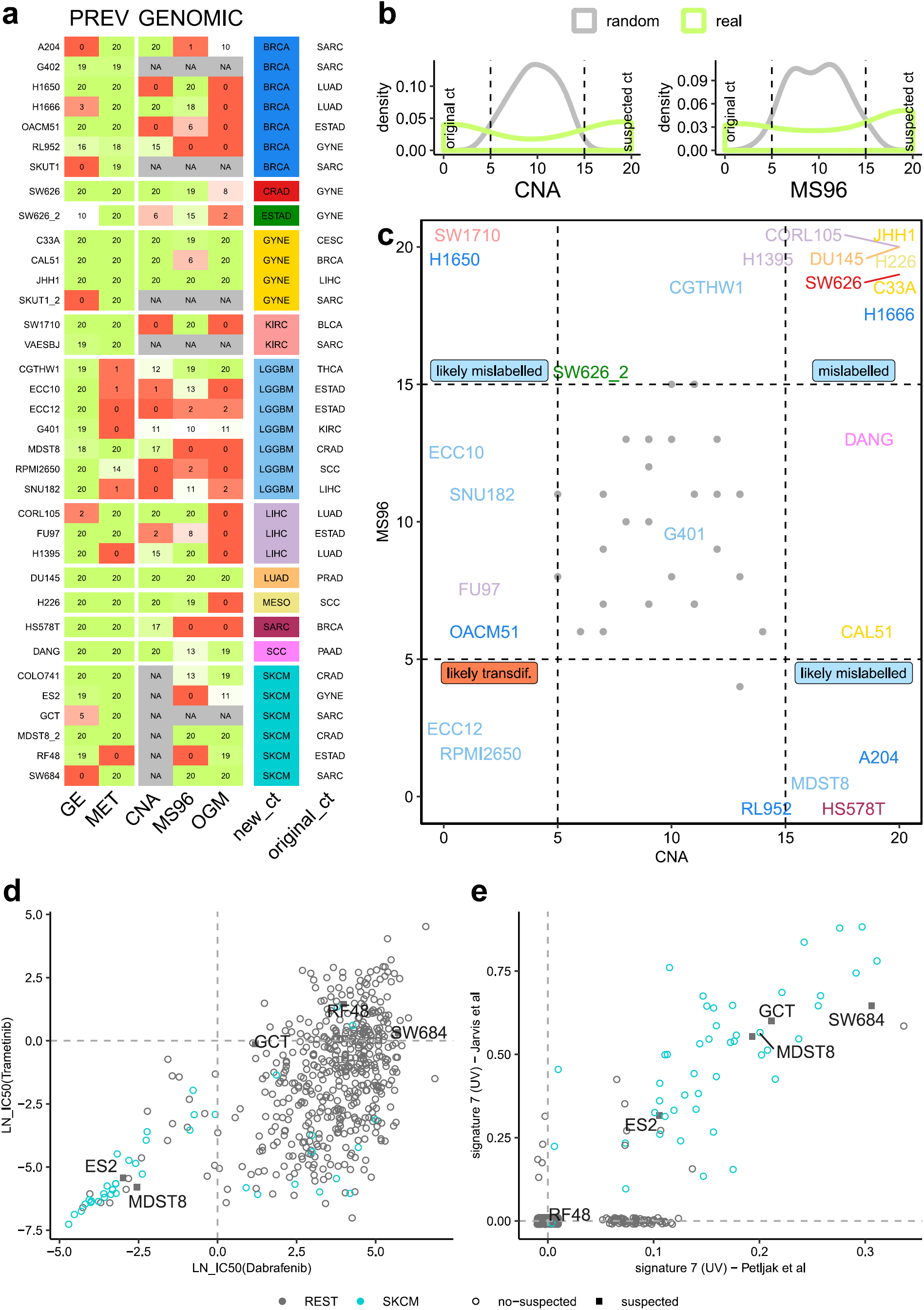
Further evidence supporting tissue identity of the suspected cell lines. **(a)** Prediction score (0-20) for each suspected cell line for 20 runs of one-versus-one classifiers that predicted suspected versus original cancer type in GE, MET, copy number alteration (CNA), mutational spectrum (MS96) and oncogenic mutations (OGM). A value of 20 means that the cell line is predicted as suspected consistently in the 20 runs of the algorithm, and a value of 0 means it is predicted as original cancer type the 20 runs. **(b)** Histograms of the prediction scores for CNA and MS96 for the models based on actual data, and a baseline on randomized data (shuffling the labels). **(c)** Prediction scores for MS96 and CNA for the suspected cell lines. Colors represent the suspected cancer type (see column “new_ct” in panel a). Grey dots represent the random values. **(d)** Drug sensitivity (IC50) for mutant BRAF inhibitors dabrafenib and trametinib for 614 cell lines. Cell lines originally labelled as skin cancer shown in red, and skin-suspected cell lines are marked with a square and their sample id. **(e)** Burden of UV-associated mutation Signature 7 (estimated from two different sources) in 614 cell lines. Cell lines originally labelled as skin cancer are shown in red and skin-suspected cell lines are marked with a square and the sample id.

Of these, approximately two-thirds (*n*=22 cell lines) received high support for the new tissue label by one or more genomic classifiers (Fig 3a; score>=15, corresponding to FDRs of 0%, 0% and 18% for the CNA, OGM and MS96 respectively, based on randomized data; Fig 3b). This data suggests 22 cell lines are candidates for assignment to another cancer type, based on converging evidence from the levels of the genome, epigenome and transcriptome, which provides confidence. Reassuringly, this list contains two cell lines which have been previously shown to be misclassified: SW626 which was initially annotated as ovarian cancer but later discovered to be derived from colon cancer (26), and COLO741 which was originally thought to be a colon adenocarcinoma cell line but later shown to originate from a melanoma (27). The fact that these two known examples were detected and reassigned to the correct cancer type provides evidence that our method is overall reliable.

The two plausible reasons why a cell line thought to originate from one cell type would need to be reassigned to a different cell type are (i) that at the time of isolation, the cell line was not of the type that it was thought to be (mislabeling), or (ii) that during prolonged cell culture, the cell line diverged greatly and now resembles another cell type (transdifferentiation). Our data allows to examine how prevalent each case is: mislabelling is expected to be reflected equally in both the epigenome and the genome, while transdifferentiation is expected to be reflected more strongly in the (presumably more malleable) epigenome, and less so in the genome, which retains the mutations from the original tumor. We suggest that mislabelling at isolation is a much more common scenario (Fig 3c, many reassigned cell lines are in the upper-right corner). However, it is possible that there exist individual examples of cell lines that have effectively transdifferentiated during culture, because their genomic features are consistent with the original tissue identity while the epigenomic features are consistent with another tissue (Fig 3c, lower left corner, e.g. the RPMI2650 and OACM51 cell lines are possible candidates).

### 4. Validation of cell lines suspected to originate from the skin

From the previous analysis, we identified a total of six cell lines which are reassigned from various cancer types to skin cancer. We note that, of skin cancers, the TCGA study contains only melanoma but not the non-melanoma skin cancers, so we are currently not able to distinguish between cell type identities of different types of skin cancer.

To further support that these cells are indeed skin cancer cells, we performed an independent analysis based on mutational signatures to confirm the mislabelling. Large-scale analyses of trinucleotide mutation spectra across human tumors have revealed at least 30 different types of mutational signatures (28). Of these, Signature 7 (C>T changes in C<u>C</u> and T<u>C</u> contexts) was associated with exposure to UV light and is highly abundant in sun-exposed melanoma tumors (29). The same signatures were recently estimated in cancer cell lines by two related methods (30,31), which enabled us to use existence UV-linked Signature 7 to examine whether these cell lines originated from the skin. Based on mutational burden of Signature 7, the known melanoma cell lines (turquoise dots) are clearly separated from the rest (Fig 3e), meaning the approach can distinguish skin-derived cells. Among the melanoma cell lines with high mutational burden of Signature 7, we found four out of five of the suspected cell lines (Fig 3e), in particular GCT, SW684, ES2 and MDST8 are very likely skin cells, and not sarcoma, sarcoma, ovarian cancer or colorectal cancer, respectively, as originally thought. For the sixth suspected cell line COLO741, the mutational signature data is not available, however COLO741 has been previously reported of being melanoma based on the expression of skin-specific genes (27).

The RF48 cell line (originally considered stomach, here putatively reassigned to skin) does not exhibit the UV signature nor the DNA methylation patterns of skin, therefore a highly confident call cannot be made. Nonetheless, a pattern of cancer driver mutations in RF48 suggests it is indeed not a stomach cell line (Fig 3a). Past work based on gene expression suggested that RF48 is indeed not representative of stomach -- instead, a lymphoid origin was proposed for RF48 (32).

Next, we sought to substantiate these findings using drug sensitivity data. In particular, two drugs (dabrafenib and trametinib) that target mutant BRAF are approved for treating melanoma in the clinic. These drugs are known to have poor efficacy in other cancer types bearing BRAF mutations, such as in colon cancers (33) and therefore sensitivity to these drugs adds confidence we are in fact looking at a melanoma cell line; (note that the converse does not necessarily hold here: resistance does not imply it is not a melanoma). Therefore, we compared the IC50 of these two drugs for all cell lines (Fig 3d). As expected, many melanoma cell lines cluster at low values of IC50 for the two drugs, meaning these cells are sensitive to the drugs. Among this cluster we observed two out of five of our suspected cell lines (ES2 and MDST8) providing further supporting evidence these are of skin, likely melanoma skin cancer origin.

In conclusion, from the six cell lines suspected of originating from skin, four of them are confirmed by the UV mutational signatures and two of those are additionally confirmed by the drug sensitivity to BRAF inhibitors. This striking example demonstrates how the transcriptome and epigenome-based tissue/cell type classifiers are able to link cultured human cell lines with their correct cancer type of origin.

In addition to these examples of skin cell lines, we have further supported several other cancer type reassignments using drug sensitivity data (34) (results summarized in Table S2). This provided evidence that the DANG cell line is consistent with squamous cell carcinoma of the lung or of the head and neck (SCC), rather than with its original assignment of pancreatic adenocarcinoma (this reassignment is also observed with multiple genomic classifiers; (Fig 3a). Similarly, SW1710 may be a kidney, rather than a bladder cell line, based on the original reassignment via transcriptome and epigenome, based on mutational patterns (Fig 3a) and additionally supported in the global analysis of drug responses (Table S2). We note that such analyses of drug screening data can be applied to distinguish only certain pairs of tissues and not all reassignments can be reliably validated in this test (see AUC scores in Table S2).

### 5. Identification of subtypes for cell lines using multi-omics analyses

Tumors are heterogeneous and major differences exist between tumor samples of the same cancer type. To manage this variability, researchers have attempted to subdivide each cancer type based on their molecular characteristics, including global patterns in gene expression and DNA methylation (35–37). However, with the exception of a few tumor types, in particular breast cancer, molecular subtypes are still being established or refined, in order to better predict disease progression in response to particular treatments.

Since drug screens and genetic screens performed in cell lines have the intent of serving as models for actual tumors, it is important to establish a method that can transfer the subtype assignments from tumors to cell lines, thereby establishing which cell line(s) are the most appropriate model for which cancer subtype.

Previously, molecular subtypes from tumors have been transferred to cell lines using different strategies. For breast cancer, cell lines subtypes have been assigned mainly using Prediction Analysis for Microarrays (PAM) analysis, which is based on a restricted set of gene expression markers (38). For colorectal cancer, the cell lines were stratified into the consensus molecular subtypes (CMS) integrating transcriptomic and genomic data (39). For renal cancer, subtypes were assigned to the cell lines using gene expression data (14). In a recent pan-cancer study, subtypes have been assigned to a set of 600 cell lines (40). It has been proposed that the cell lines do not usually represent all subtypes of a particular cancer type, possibly due to a bias introduced during the process of immortalization (38,40).

Our approach to assign subtypes to cell lines herein is to apply the same strategies that have allowed us to get accurate cancer type classifiers: first, the integration of transcriptomic and epigenomic data to boost confidence in the predictions, and second, careful adjustment of the two data types to make them comparable between TCGA tumors and cell lines (Fig S1).

An important consideration in the task of inferring the cell lines’ subtypes is the absence of true labels needed for systematic validation, thus assignments should be treated as tentative. However, for breast cancer cell lines the subtype labels are available (38) and can be used as a benchmark of our multi-omics based methodology.

We examined proposed subtypes for 15 cancer types in TCGA and generated subtype classifiers (Methods) for each cancer type. The combination of both data types (GE and MET) achieved a higher cross-validation accuracy in the TCGA (median AUPRC across cancer types: 0.81) than GE (0.76) or MET (0.72) separately. Therefore, we used the combined datasets to generate subtypes classifiers and propose assignments of the cell lines to cancer subtypes. Since we are using one-*versus*-rest classifiers each cell line can be assigned to more than one subtype. However, the majority of them are assigned to only one subtype (Fig S6a); we used only those in further analysis. As a benchmark, we calculated the accuracy for the breast cancer cell lines with subtypes available (Fig S6b): the median AUPRC (across breast cancer subtypes) for CL is 0.83. This suggests acceptable performance in obtaining tentative subtype assignments for cell lines in all 15 cancer types, which we provide as a resource in Table S3. This resource is complementary to a recent set of subtype predictions for 9 cancer types based on transcriptomes (40).

Next, we examined if the relative prevalence of subtypes is similar between tumors and cell line panels of the same cancer type. Cell line panels of some cancer types have good representation of subtypes, for instance lung squamous cell cancer, head and neck squamous cell cancer, lung adenocarcinoma, and gastric/esophageal cancers (Fig S6c). However, the converse is the case for liver, skin and thyroid cancer cell lines, in which a single subtype predominates in cell line panels but not in tumors (statistics listed in Supplementary Table 4). Additionally, we observe suboptimal representation (where half of the tumor subtypes are not represented) in the kidney, bladder and brain cancer cell line panels, when considering the 463 cell lines we analyzed. This suggests that -- in some cancer types more than others -- the commonly used cell line panels do not represent the diversity of molecular subtypes in tumors, which should be taken into account when interpreting experimental data. One possible reason is the relative ease of culturing certain subtypes, compared to others (2).

### 6. Accounting for mislabelled cell lines reveals new associations in drug screening data

We detected 35 cell lines that may have a tissue or cell type identity different than the one originally assigned to them. Because the cell type is an important determinant of drug response in cancer cell lines and in tumors (11), we hypothesized that the inclusion of this new tissue information into analyses of genetic determinants of drug sensitivity may change the results. In a comprehensive study, Iorio *et al.* searched for associations between drug response and Cancer Functional Events (CFEs): the recurrent mutations, CNA and hypermethylation events present in human tumors (11). Here, we used GDCSTools (27) to reproduce the results of that study, however after filtering the cell lines to those that better represent the cancer type in question. In particular, we repeated the same analysis using for each tissue (i) all the cell lines; (ii) only the cell lines in the ‘golden set’ (G); (iii) as a less stringent filtering criterion, only the cell lines in the ‘golden and silver set’ (G&S). Additionally, as controls we included a random subset of cell lines that matches (iv) the number of cell lines in ‘golden set’ (r_G) and (v) the number in ‘golden and silver set’ combined (r_G&S).

For the majority of the cancer types, we observed that one of the filtered subsets recovered a higher number of significant (at FDR<= 25%) associations of CFE with drug sensitivity or resistance, than were recovered using all cell lines (Fig S7). For instance, for glioblastoma, using the ‘golden set’ cell lines we found 23 new associations, which were not recovered from the entire cell line panel nor from the random-subset controls (Fig 4b). For example, this recovers the positive association of CDKN2A loss with camptothecin sensitivity (Fig 4c), which was previously reported in an independent analysis of the NCI-60 cell line panel screening data (41). Similarly, for pancreatic adenocarcinoma, benefits were observed by focusing on cell lines that resemble the corresponding cancer type better: using only the ‘golden set’ plus ‘silver set’ cell lines, 10 new significant associations were found (Fig 4b). For instance, we detected that SMAD4-mutant cell lines are more resistant to piperlongumine, a natural product claimed to have antitumor properties exerted via multiple pathways (42,43). Mutations of the tumor suppressor gene EP300 were associated with higher sensitivity to three drugs in pancreatic cancer cell lines (Fig 4d).

**Fig. 4.**
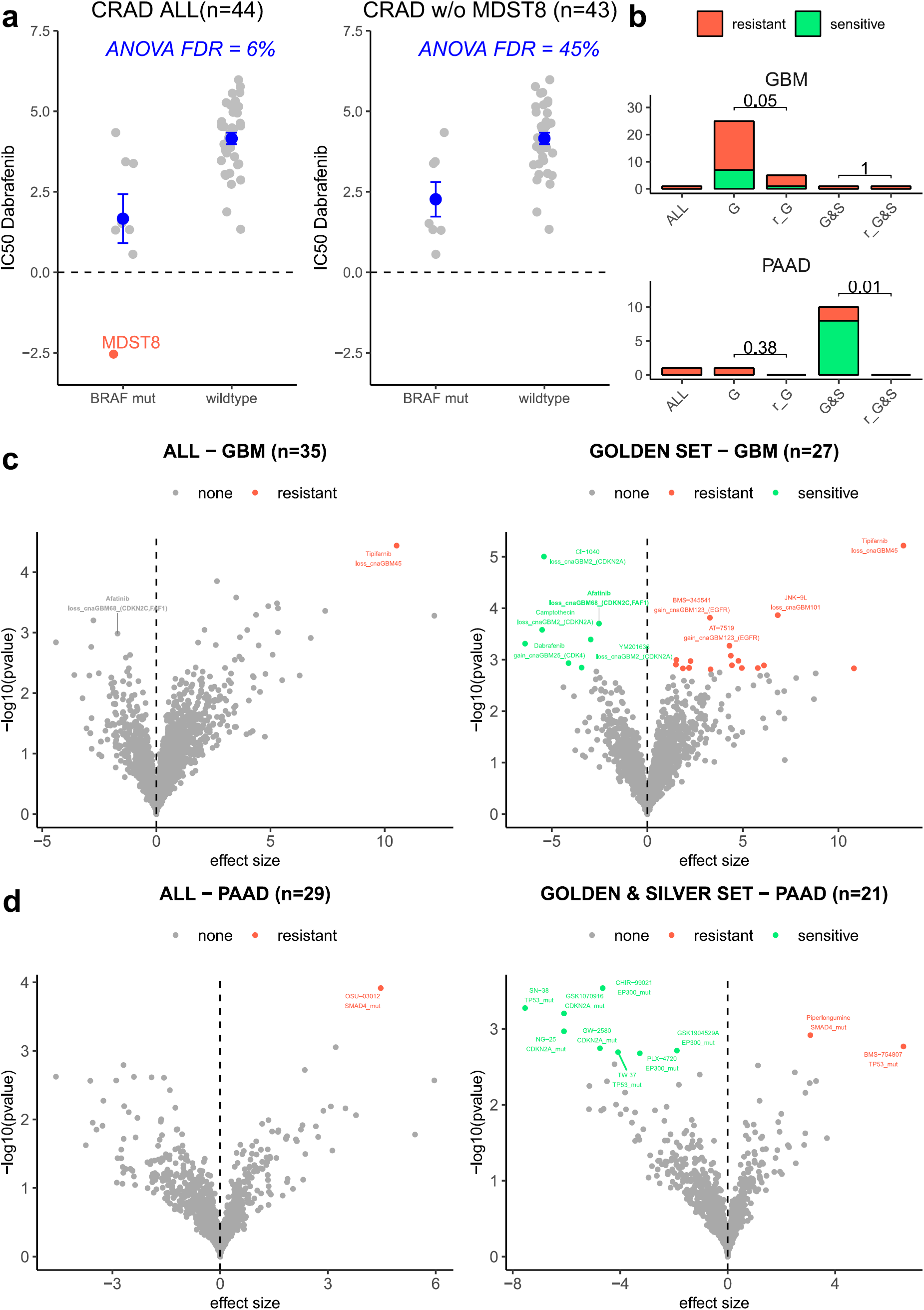
Drug sensitivity association testing using high-confidence sets of cell lines. **(a)** Drug sensitivity (IC50) to dabrafenib in all CRAD cell lines (left) and all CRAD cell lines except MDST8, which is suspected of being skin cancer (right). Two groups are compared: cell lines with BRAF mutation and without (wild-type). ANOVA FDR for this association (dabrafenib and BRAF mutation) shown in blue for both datasets. Horizontal line is shown at 0, because score <0 implies sensitivity to the drug. **(b)** Number of significant associations between Cancer Functional Events (CFEs) and drugs detected in the ANOVA test for all cell lines (ALL), cell lines in the golden set (G), cell lines in the golden plus silver set (G&S), random subset of cell lines that match the number of cell lines in the golden set (r_G) and in the golden plus silver set (r_G&S). For the random subsets, the number of significant associations is calculated 10 times (with different random selection) and the median of the 10 runs is shown. P-values for a sign test (one-tailed, alternative = “less”) between the number of associations in the G/G&S versus the number of associations in r_G/r_G&S are shown. See Fig S7 for remaining cancer types. **(c)** Differential sensitivity of drugs were analysed by ANOVA for all brain cancer cell lines (left) and the brain cancer cell lines in the golden set only (right). Each point is an association between the sensitivity of a drug and a genetic feature (CFE). **(d)** Differential sensitivity of drugs were analysed by ANOVA for all pancreatic (PAAD) cell lines (left) and PAAD cell lines in the golden and silver set only (right). Each point is an association between the sensitivity of a drug and a genetic feature (CFE).

The observation that more associations were found despite using a somewhat lower number of cell lines (thus less statistical power) emphasizes the importance of using the cell lines that more closely model the tissue and/or cell type-of-origin of the cognate tumor.

Colorectal cancers provides an illustrative example of how important is to remove nonrepresenative cell lines from drug screening efforts. In the Iorio *et al.* (2016) study, 50 colorectal adenocarcinoma (CRAD) cell lines were tested. Of those, we strongly suspected that MDST8 derives from skin. To test the influence of this individual mislabelled cell line we have performed association testing with all CRAD cell lines, and after excluding MDST8. For the association of the drug dabrafenib with BRAF mutation status, we observed that all CRAD cell lines (irrespective of BRAF mutation) are not sensitive, except for MDST8 which is strongly sensitive (Fig 4a). The FDR of the ANOVA analysis when using all cell lines is 6%, while when removing MDST8 the FDR is 45%. Therefore, in this case, the presence of a single mislabelled cell line is sufficient to cause the appearance of a false association between a drug and a feature. This is fully consistent with clinical responses: in contrast to the good response of patients with BRAF-mutant melanoma to dabrafenib, colorectal tumors with the same BRAF V600 mutation are not sensitive to BRAF or MEK inhibitor monotherapy (33).

### 7. Accounting for mislabelled cell lines reveals new associations in genetic screening data

Motivated by the many novel associations revealed by reanalyzing the drug screening data, we asked if the same extends to genetic screening data in cancer cell lines, because results in genetic screens may also depend on cell lineage (12). To further investigate, we analyzed CRISPR screening data from Project Score and Project Achilles (see Methods), from which 347 cell lines overlap our tested cell lines. Then, we applied the same association testing method, which was however underpowered because the number of available overlapping cell lines was smaller. Nonetheless, in colorectal and ovarian cancer, we observed that focussing only on the ‘golden set’ and/or ‘silver set’, the number of associations recovered increased (as a control, there were no increases in the random cell line subsets of the same size; Fig 5a, Fig S8).

**Fig. 5.**
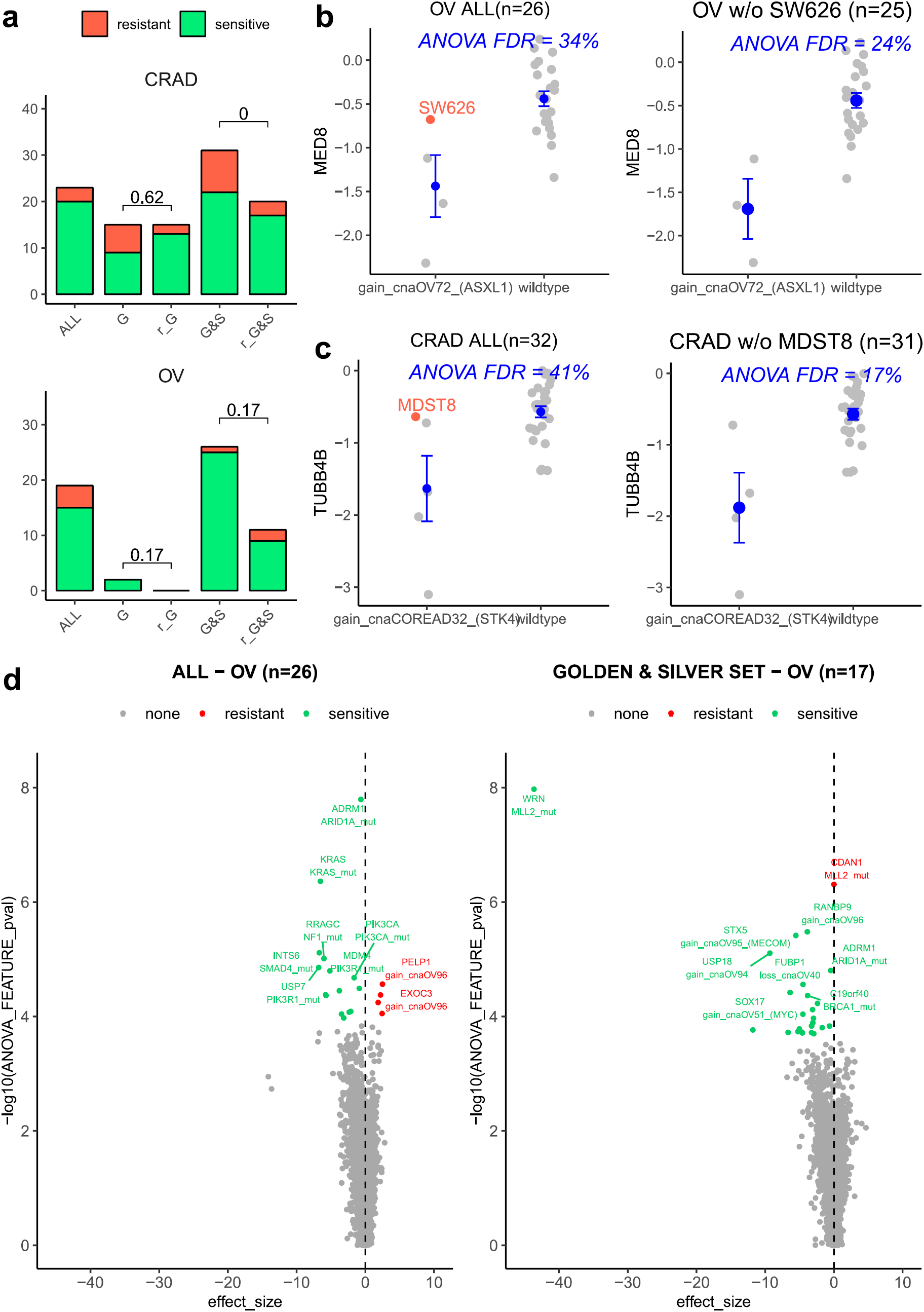
Analysis of genetic screening data using high-confidence cell lines. **(a)** Number of significant associations between Cancer Functional Events (CFEs) and gene dependencies (in CRISPR knockout screens) detected in the ANOVA test for all cell lines (ALL), cell lines in the golden set (G), cell lines in the golden and silver set (G&S), random subset of cell lines that match the number of cell lines in the golden set (r_G) and in the golden and silver set (r_G&S). For the random subsets, the number of significant associations is calculated 10 times and median. P-value for a sign test (one-tailed) between the associations in the G/G&S and the associations in the 10 runs of r_G/r_G&S are shown. See Fig S8 for remaining cancer types. **(b)** Fitness effect (fold change) for MED8 k.o. in all OV cell lines (left) and all OV cell lines except SW626, which is suspected of originating from CRAD (right). Two groups are compared: cell lines with copy number gain in a region containing ASXL1 (gain_cnaOV72), and without (wild-type). ANOVA FDR for this association (MED8 k.o. and gain_cnaOV72) is shown in blue for both datasets. **(c)** Fitness effect (fold change) for TUBB4B k.o. in all CRAD cell lines (left) and all CRAD cell lines except MDST8, which is suspected to originate from skin (right). Two groups are compared: cell lines with copy number gain in region containing STK4 (gain_cnaCOREAD32) and without (wild-type). ANOVA FDR for this association (TUBB4B k.o. and gain_cnaCOREAD32) shown in blue for both datasets. **(d)** Differentia ldependency biomarkers were analysed by ANOVA for all ovarian cancer (OV) cell lines (left) and OV cell lines in the golden and silver set only (right). Each point is an association between the fitness effect of a gene and a genetic feature (CFE).

To illustrate the importance of removing suspect cell lines in gene dependency screenings, we provide two examples of associations that were originally not detected as significant due to the presence of a mislabelled cell line. For ovarian cancer, the presence of SW626 (mislabelled cell line confirmed by the literature (26)) prevents finding the association between MED8 dependency and a copy number gain in the region containing ASXL1 (cnaOV72) as significant (Fig 5b). Similarly, for colorectal cancer the presence of MDST8 (mislabelled cell line confirmed by the UV mutational signature) prevents finding the association between TUBB4B dependency and a copy number gain in STK4 (cnaCOREAD32) (Fig 5c). Finally, a significant association between WRN dependency and MLL2 (also known as KMT2D) gene mutation is recovered only with the filtered cell lines in ovarian cancer (Fig 5d). This WRN-MLL2 association has been recently reported using a somewhat different set of cell lines (from Project Score) (44) that partially overlap our set.

Finally, our re-analyses of drug screening and genetic screening data revealed an interesting association independently supported in both drug and genetic data. The drug afatinib inhibits the EGFR protein and is clinically indicated for EGFR-mutated lung cancer, however in EGFR-altered glioblastoma afatinib is generally not considered to elicit a response (45). Consistently, afatinib sensitivity was associated with EGFR alterations in lung cancer previously (11), as well as in our re-analysis (FDR G&S = 0.6%), but not in the brain cell line panel (all FDR ≥ 25%). However, using the focussed (golden set) of brain cancer cell lines revealed a significant association (ANOVA FDR = 15%, Fig S9a) between afatinib sensitivity and a different genetic alteration: copy number loss in a region at 1p32.3 containing the CDKN2C and FAF1 genes (id: cnaGBM68). Remarkably, the same loss at 1p32.3 is associated with sensitivity to genetic knockout of EGFR in brain cell line panels in two independent large-scale genetic screens (Project Scores and Project Achilles, Fig S9cd) and another drug screen (PRISM, Fig S9b). The meta-analysis of the two drug screens and two genetic screens suggests high strength of combined evidence (p=0.00094, Fisher’s method of combining p-values) linking the loss at 1p32.3 (chr1: 51169045-51472178) with sensitivity to pharmacological or genetic EGFR inhibition in brain cells, suggesting a strong candidate for follow-up work.

In summary, the presence of cell lines with dubious or incorrect labels of tissue identity may strongly impact association studies of drug or CRISPR screening data in two different ways. First, the presence of mislabelled cell lines can cause the appearance of spurious associations that do not reflect the biology of the cancer type of interest. Second, the presence of mislabelled or divergent cell lines can prevent the recovery of true associations.

## Discussion

Cell lines are commonly used as models for tumors, however it is an open question how to best apply the available cell line panels to learn about cancer biology. The availability of genomic data from large tumor cohorts and from cell line panels has spurred multiple efforts to find which cell line(s) are closer to tumors by their transcriptomic (10,39,40,46) and/or genomic features (13,14), presumably making better models, and which are more distant from examples of actual tumors, presumably making less good models of tumor biology.

Our work addresses a different question: we attempt to detect the cancer type (i.e. tissue and/or cell type) that originated the cell line, in order to ascertain if this matches the declared origin of the cell line. A mismatch may conceivably stem from the sampling step, for instance a metastasis might have a different tissue-of-origin than thought at time collection. The work-up after collecting the tumor sample may have inaccurately assigned the cancer type based on unclear histological or anatomical features. Another possibility is that the mismatch might stem from the step of adaptation to cell culture, where a minority cell type not representative of the tumor prevails over other tumoral cells. We consider these to be cases of cell line mislabeling during isolation. In addition, we would also detect cases where the cell line might have acquired some properties of a different tissue/cell type during culture, however our analyses (Fig 3c) suggest this is a less common occurrence, although individual examples of this cannot be ruled out.

Importantly, this phenomenon of tissue / cell type mislabeling is distinct from well-known and widespread cell line misidentification issues (23,47), where one cell line (often HeLa) was mistakenly used in place of another cell line, commonly due to cross-contamination. The cell line panels that provided data used in our analyses (GDSC and CCLE) have authenticated their cell lines (4,44), thus misidentification/cross-contamination cannot underlie our observation that the mislabeling of the cancer type of origin is not uncommon. (We note there were rare cases of misidentified cells reported in these panels (44) however these do not overlap our results.)

Methodologically improving over previous work, we introduce the HyperTracker framework that performs global analyses that independently examine transcriptomic, epigenomic and several types of mutational features. Additionally, while carefully adjusting for the known bulk differences between cell lines and tumors, which might have resulted e.g. from impurities in tumors, or from altered expression of cell-cycle-related genes in cell lines (40,48). Parallel analyses of different omics data provides increased confidence in our inferences, which suggested, remarkably, that 5.7% (35 of total 614 considered cell lines) exhibit significant transcriptomic and epigenomic features of a different tissue/cell type than the declared cell type of origin. For 3.6% (22 cell lines) these reassignments to a different cancer type were additionally supported in at least one type of genomic evidence. This increased confidence that these were indeed examples of cell lines with mislabeled (or, less likely, diverged) tissue/cell-type identity. A striking example are cell lines GCT, SW684, ES2 and MDST8 that we predict to originate from the skin, based on the presence of the UV mutational signature, in addition to transcriptome/epigenome data. These cases are reminiscent of the recent reports of UV mutational signatures found in some cases of presumed lung cancers, suggesting they may instead be metastases originating from a sun-exposed area of skin (49).

In interpreting our data, an important consideration is that the cancer samples types in TCGA may not necessarily reflect the full diversity of rarer subtypes within a cancer type, which may cause some ambiguous predictions. For instance, the ECC10 and ECC12 cell lines are assigned to STAD cancer type (stomach adenocarcinoma) when matched with TCGA tumors. However, these cell lines originate from gastric small cell neuroendocrine carcinomas. This may explain why, in our analysis, gene expression patterns point towards brain tissues, while mutational features suggest stomach cancer. In such cases of disagreement between different types of features, a future use of a more exhaustive set of reference tumor data may help resolve the ambiguity and improve confidence in predictions.

The genomic classifiers we employed here were based on whole-exome sequences and were overall less powerful than the transcriptome/DNA methylation classifiers in our data (Fig S4). Recent work by us and others (24,25) suggests that analyzing whole-genome sequences of these cell lines would permit use of additional, highly predictive features based on regional mutation density of chromosomal domains. This may provide further genomic evidence for the identity of the cell-of-origin for the 35 suspected cell lines. Experimental work on these cell lines will provide further evidence to support or refute our predictions based on global analyses of omics data.

Knowing the correct tissue-of-origin label for a cell line is important, because this has a strong bearing on the response of the cell line to drug treatment and to genetic perturbation. We demonstrate the implications of this general principle to analyses of drug and genetic screening data: by accounting for suspect cell lines, the power to discover new determinants of sensitivity to drug/genetic perturbation may increase substantially for some cancer types, such as brain, lung and pancreatic cancers. Therefore, when designing future screening efforts, it is not only important to increase the number of cell lines to gain more power, but it is also important to focus on the cell lines that are most consistent with the tissue and/or cell type of interest.

## Methods

### Omics data collection and preparation

#### DNA methylation data

We downloaded DNA methylation data as beta values (platform Illumina Human Methylation 450) from GDC Data Portal (50) for TCGA samples and from Genomics of Drug Sensitivity in Cancer (GDSC) (3) for CL samples. We filtered out all probes outside promoter regions and probes with NA values in more than 100 samples. For the probes in promoter regions, we selected only one probe per gene, keeping the probe with the highest standard deviation (sd) across samples. We transformed the beta-values to m-values (log2 ratio of the intensities of methylated probe versus unmethylated probe). In total, we have 10,141 features for 942 CL samples and 8,453 TCGA samples.

#### Gene expression data

We downloaded gene expression data as transcripts per million (TPM) from GDC Data Portal (50) for TCGA samples and from GDSC (3) and the Cancer Cell Line Encyclopedia (CCLE) (4) for CL samples. We filtered out genes with NA values in more than 100 samples and selected the overlapping genes between the 3 sources. We removed low expressed genes (TMP<1 in 90% of the samples). We applied square root transformation to the data. In total, we have 12,419 features for 942 CL samples and 9,149 TCGA samples.

Finally, for both DNA methylation (MET) and gene expression (GE), we created datasets of different sizes: 1,000; 2,000; 3,000; 5,000; and 8,000 features by selecting the genes/probes with the highest standard deviation across TCGA samples only.

#### Copy Number Alteration data

We downloaded Copy Number Alteration data (computed by gene) from GDC Data Portal (50) for TCGA samples and from DepMap (51) for CL samples. In total, we have 20,491 features for 942 CL samples and 9,188 TCGA samples. To reduce the dataset, we selected 299 cancer driver genes (52) and filtered out the rest.

#### Mutation data

For human tumors, we downloaded mutation data as whole exome sequencing (WES) MC3 dataset (53) from the GDC Data Portal for TCGA samples. For cell lines, bam files were obtained from European Genome-phenome Archive (EGA) (ID number: EGAD00001001039). Variant calling was performed using Strelka (version 2.8.4) with default parameters. Variant annotation was performed using ANNOVAR (version 2017-07-16). In samples where Strelka was unable to run, a re-alignment was performed using Picard tools (version 2.18.7) to convert the bams to FASTQ and, following that, the alignment was perfomed by executing bwa sampe (version 0.7.16a) with default parameters. The resulting bam files were sorted and indexed using Picard tools. To account for germline variants, we removed all mutations that were present in the gnomAD database (54) at an allele frequency >= 0.001 in any of the populations. Finally, using the filtered somatic mutations we calculated three set of mutational features: Regional Mutation Density (RMD), Mutation Spectra (MS96) and Oncogenic Mutations (OGM) as described in Salvadores et. al (24). RMD features did not exhibit high accuracy when applied to exome-sequencing data and so were not considered further in this analysis.

For the cell line samples, we matched their cancer types to the TCGA cancer types using the cell line metadata from GDSC (3) and manually annotated those that did not have TCGA label using cellosaurus (55). Next, we selected the cell lines from solid tumor that had a matching cancer type in TCGA, ending up with a total of 614 cell lines from 22 cancer types. Blood cancers (LAML and DLBC) are not tested because they are commonly growth in suspension, therefore their confusion with solid tumors is less likely to occur. For further analysis, we merged the cancer types that were overall similar: HNSC with LUSC and ESCC (SCC), GBM with LGG (LGGBM), STAD with ESAD (ESTAD) and OV with UCEC (GYNE).

The identification of the cell line samples were performed by the databases providing the data using short tandem repeat (STR) analysis (4,44). Of note, they reported a few commonly misidentified cell lines: Ca9-22, RIKEN, MKN28, KP-1N, OVMIU and SK-MG-1 (44). These cell lines do not overlap with our suspected samples and additionally the misidentification does not impact tissue or cancer type of origin.

### Data alignment between tumors and cell lines

For the alignment of TCGA and CL data we first applied quantile normalization (*R package preprocessCore 1.46.0*) and second applied ComBat (*R package sva 3.32.1*), a batch effect correction method. We used ComBat as if our dataset was the TCGA and CL data combined, and the batch effects were whether a sample belongs to TCGA or CL (for MET) or a sample belongs to TCGA, GDSC or CLLE (for GE). We applied this method for GE, MET, CNA, MS96 and RMD. For validation, we calculated a principal component analysis (PCA) subsampling TCGA data to match the CL samples (stratified by cancer types). Additionally, we calculated Elastic Net (EN) classifiers to predict (in the processed dataset) TCGA *versus* CL and calculated the AUC and AUPRC to check whether the process of alignment is being successful or not.

In addition to the chosen adjustment method, we tested other approaches based on Canonical Correlation Analysis, Partial Least Squares and principal component analysis, which did not exceed accuracy of ComBat (data not shown) and therefore were not examined further.

### Cancer type classifiers

For the TCGA dataset we generated Ridge regression model for predicting the cancer type in a One-vs-Rest manner (*using cv.glmnet function with alpha=0 and family = binomial, R package glmnet 2.0.18*). To calculate the accuracy, we trained classifiers in the TCGA dataset and tested in the CL dataset. In particular, we calculated the Area Under the Receiver Operating Characteristic curves (AUC) and the Area Under the Precision Recall curve (AUPRC) for each cancer type vs the rest (all the rest of cancer types combined).

#### FDR Score

For each cell line, we calculated an FDR score of belonging to a particular cancer type. For this, we divided the TCGA data into two datasets (training and testing) of the same size keeping the cancer type proportions. For each cancer type, we trained classifiers in the TCGA training dataset and we introduced the cell lines one by one with the testing data and calculated the precision recall (PR) curve (TCGA testing + 1CL). We set the cell line FDR score for that specific cancer type as (1 - precision) at the threshold where the cell line is situated in the PR curve. Overall, for every cell line we obtained 17 FDR scores, 1 for each possible cancer type. We repeated this procedure 5 times and calculated the median FDR for every case to get more robust values. In addition, when training for 1 cancer type (label = 1) versus the rest of cancer types combined (label = 0) we made some exceptions and removed those cancer types which are similar and therefore the classifier is not good at separating them (e.g. when we calculated FDR for ESTAD, we removed from the rest CRAD and PAAD, all hidden cases in Table S7). This is conservative with respect to reassigning cell lines to another cancer type, however some resolution is traded off because the more closely related cancer types are, by design, not distinguished. We have further attempted to reclassify cell lines within these hidden tissues and the combined ones. However, when using One-vs-One classifiers the accuracy is not good enough for distinguishing the two cancer types in the cell lines (data not shown).

Once we have a list of suspected cell lines, we have an “original” cancer type and a “suspected” cancer type. Therefore, we generated One-vs-One classifiers (original *versus* suspected) using TCGA dataset (balancing the classes) and for each suspected cell line we checked if it is predicted as “original” or “suspected”. We repeated this prediction 20 times and counted the number of times a cell line is predicted as suspected. Therefore, we defined a prediction score (range between 0 and 20) for every cell line, where 0 means never predicted as suspected and 20 always predicted as suspected. As a control, we repeated the same procedure with randomized cancer type labels. We calculated this prediction score for GE, MET, CNA, OGM and MS96 datasets. For calculating the FDR at a score ≥15, we applied this formula: FDR = FP/(FP+TP) where FP was the number of cell lines with score ≥15 in the randomized data and TP the number of cell lines with score ≥15 in the actual data.

### Independent validation

We downloaded drug sensitivity for the CL from GDSC (3). From all the drugs we selected *trametinib* and *dabrafenib*, FDA-approved drugs for melanoma treatment. We compared IC50 values for these two drugs for all cancer types.

We downloaded mutational signatures from cell lines available in Jarvis *et al* (31) and Petljak *et al* (30) and we compared the exposures of all cell lines for Signature 7 (UV light). In Petljak *et al* dataset the signature 7 is divided into Signature 7a, b, c and d. Therefore, we used the sum of exposures across all four subtypes of Signature 7.

We downloaded another set of drug screening data (PRISM 19Q3) (34) for the CL dataset. For the suspected cell lines, we generated One-vs-One classifiers (*using cv.glmnet function with alpha=1 and family = binomial, R package glmnet 2.0.18*) for predicting original *versus* suspected cancer type based on the drug sensitivity data. We performed 20 runs of each case and count how many times it is predicted as suspected (prediction score 0-20). Additionally, we calculated the AUC for each classifier.

### Subtype classifiers

We downloaded cancer subtypes for the TCGA samples from the *R package TCGAbiolinks 2.12.6* (56). We combined the GE and MET datasets. For this data, we generated Ridge regression model for predicting the subtypes in a One-vs-Rest manner (*using cv.glmnet function with alpha=0 and family = binomial, R package glmnet 2.0.18*) within each cancer type. We trained models in TCGA and we predicted subtypes for the cell lines. Additionally, we used cell line’s subtypes for breast cancer from a previous paper (38) to calculate the confusion matrix and the AUPRC.

We performed a chi-square test (*R package stats 3.6.0*) and calculated the cramer’s V statistic (*R package lsr 0.5*) for checking whether the proportion of subtypes between TCGA and CL is maintained for each cancer type.

### Drug and CRISPR screening data

We downloaded drug sensitivity and cancer funcional event (CFE) data from the Iorio *et al*. study (11). Cancer Functional Events (CFEs) are a collection of recurrent mutations, CNA and hypermethylation events present in human tumors (11). We used GDSCTools (57) to search for associations between the drugs and the CFEs in every cancer as they did. In particular, we performed this analysis using for each tissue (i) all the cell lines; (ii) only the cell lines in the golden set (G); (iii) only the cell lines in the golden and silver set (G&S). Additionally, as controls we included a random subset of all cell lines matching (iv) the number of cell lines in goldenset (r_G) and (v) the number in golden and silver set combined (r_G&S). We counted the number of significant hits (FDR≤25%) for each of the cancer types. For the controls, we repeated the subsampling 10 times and took the median of significant hits. We compared the number of hits for all the cell lines (same as in Iorio *et al* study) with the number of hits for the different subsets of cell lines according to our grouping. Additionally, we performed a sign test (*R package BSDA 1.2.0*) comparing the significant hits in the G/G&S subsets versus the significant hits over 10 runs in the random G/ random G&S and calculated the p-value for all cancer types (alternative = “less”).

Similarly, we downloaded gene dependency data from Project Score (44) and Project Achilles (58) processed with the Project Score pipeline and combined them. From a total of uniquely 696 cell lines, 357 overlap with the 600 cell lines tested with our method. For those 357 tested cell lines, we repeated the same procedure as described above for the drug sensitivity data.

