## Supplementary material for "Matching cell lines with cancer type and subtype of origin via mutational, epigenomic and transcriptomic patterns": Supplemenary Figures S1-S2 and S4-S9

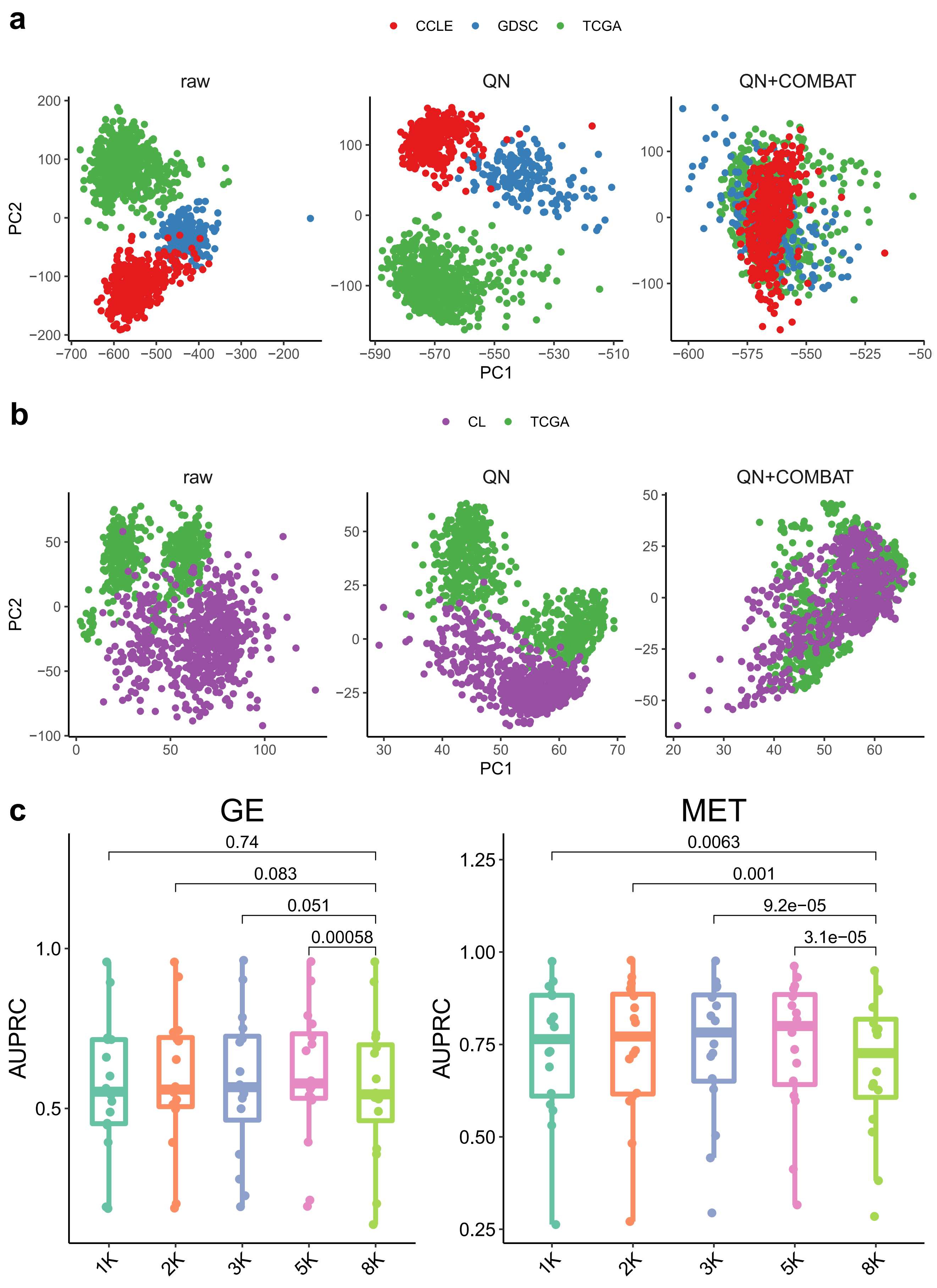


***Fig S1.*** ***(a)*** *Principal component (PC) 1 and PC2 of a PC analysis in the GE pre-adjustment data (raw), intermediate step (quantile normalization - QN) and in the post-adjustment data (QN+COMBAT). Colors represent the batch effects we are correcting for (GDSC and CCLE are two sources for the cell lines and TCGA is the source for the human tumors).* ***(b)*** *PC1 and PC2 of a PCA in the MET pre-adjustment data (raw), intermediate step (QN) and in the post-adjustment data (QN+COMBAT). Colors represent the batch effects we are correcting for (CL is the source for the cell lines and TCGA is the source for the human tumors).* ***(c)*** *Area Under the Precision Recall curve (AUPRC) for predicting cancer type (training set human tumors and testing set cell lines) in GE (left) and MET (right) at different dataset sizes (1K to 8K features). Features selected using the standard deviation across samples of the feature (gene/probe) in the human tumors. P-values for the wilcoxon-test (paired) between 8K and the rest are shown.*

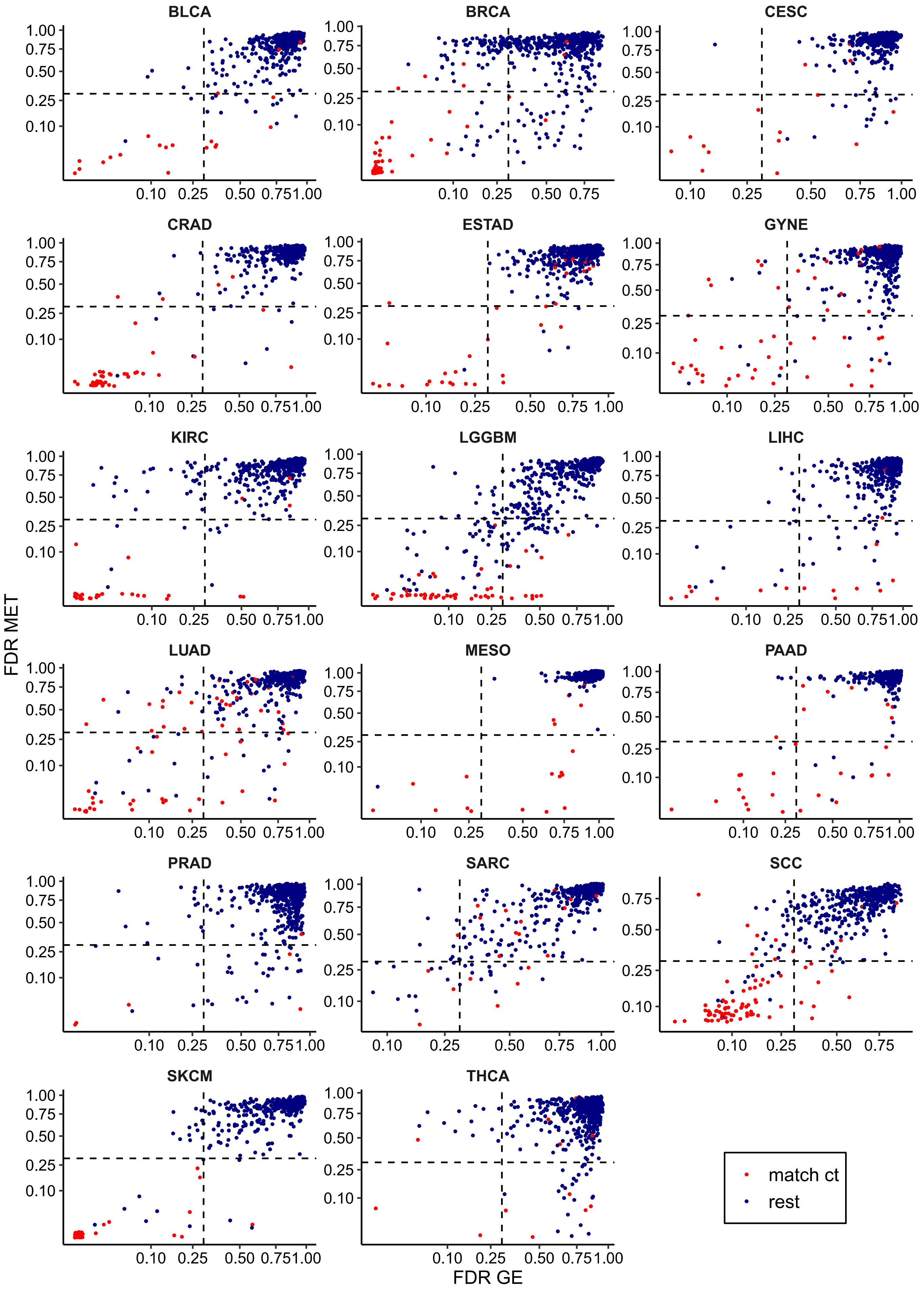


***Fig S2.*** *False Discovery Rate (FDR) scores for 614 cell lines calculated in MET and GE cancer-type classifiers (one-vs-rest scheme). The lower the FDR the more likely of belonging to that particular cancer type (shown on top of each plot). The cell lines that are originally annotated as the cancer type that is being tested are shown in red, the rest in blue.*

*( FIGURE S3 IS PROVIDED AS A SEPARATE PDF FILE )*

***Fig S3****. Heatmaps for the 25 genes (in GE) and probes (in MET) whose ridge regression coefficients presented the highest absolute values for a particular cancer type versus the rest. There is one heatmap per cancer type that present at least 1 suspected cell line (suspected cancer type written in the title). In each heatmap, the suspected cell lines from belonging to that particular cancer type are labeled on the right side. In each case, the cancer types included are the suspected one and the original cancer types from the suspected cell lines.*

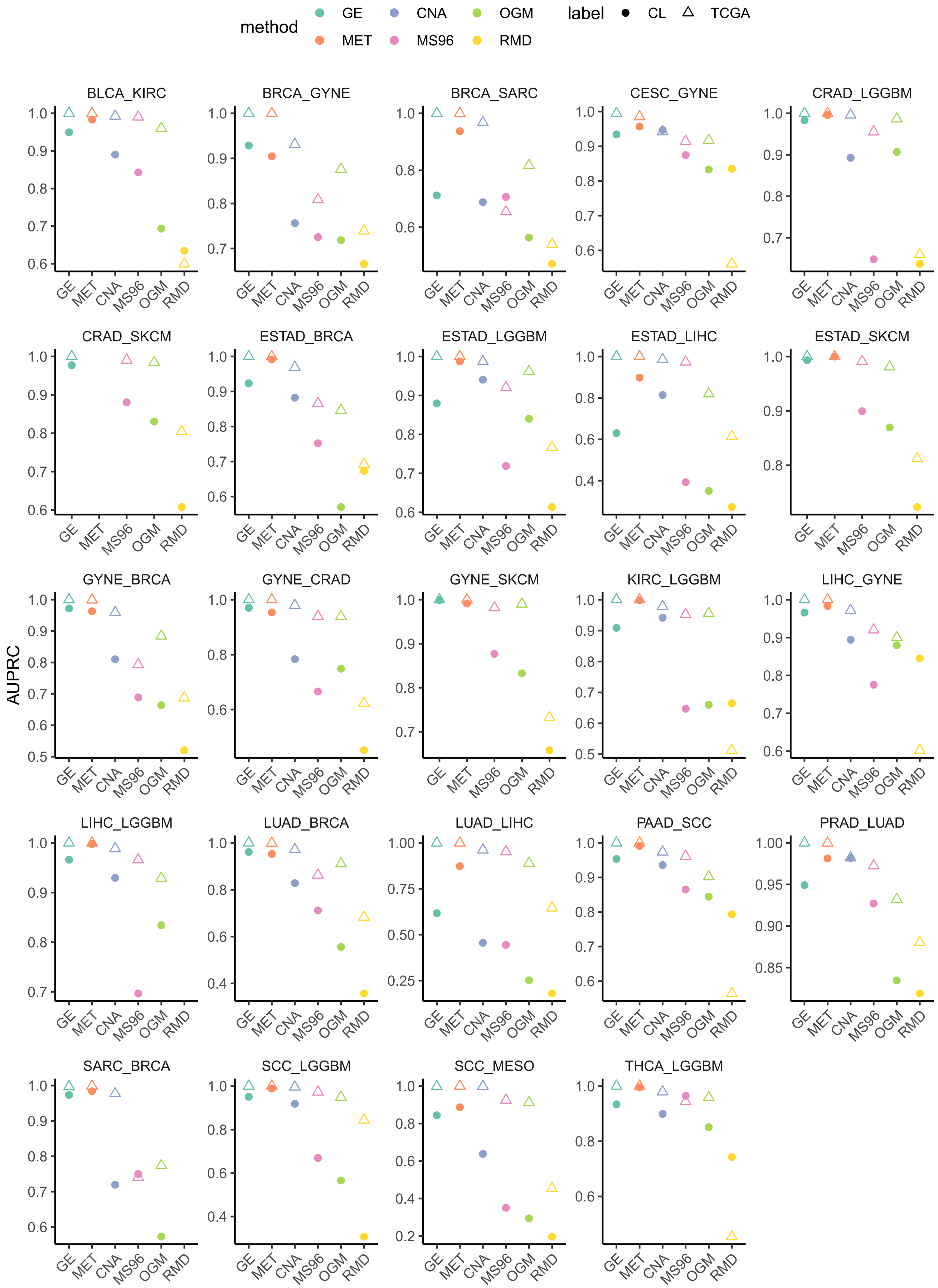


***Fig S4.*** *Area Under the Precision Recall curve (AUPRC) for predicting cancer type in one-vs-one classifiers (the two tested cancer types are shown in the top of the plot) in GE, MET, CNA, MS96, OGM and RMD. The classifier is trained in a training set of human tumors (TCGA) and the AUPRC is calculated in a testing set of human tumors (TCGA, shown with a triangle) and in all the cell lines (CL, shown with a circle).*

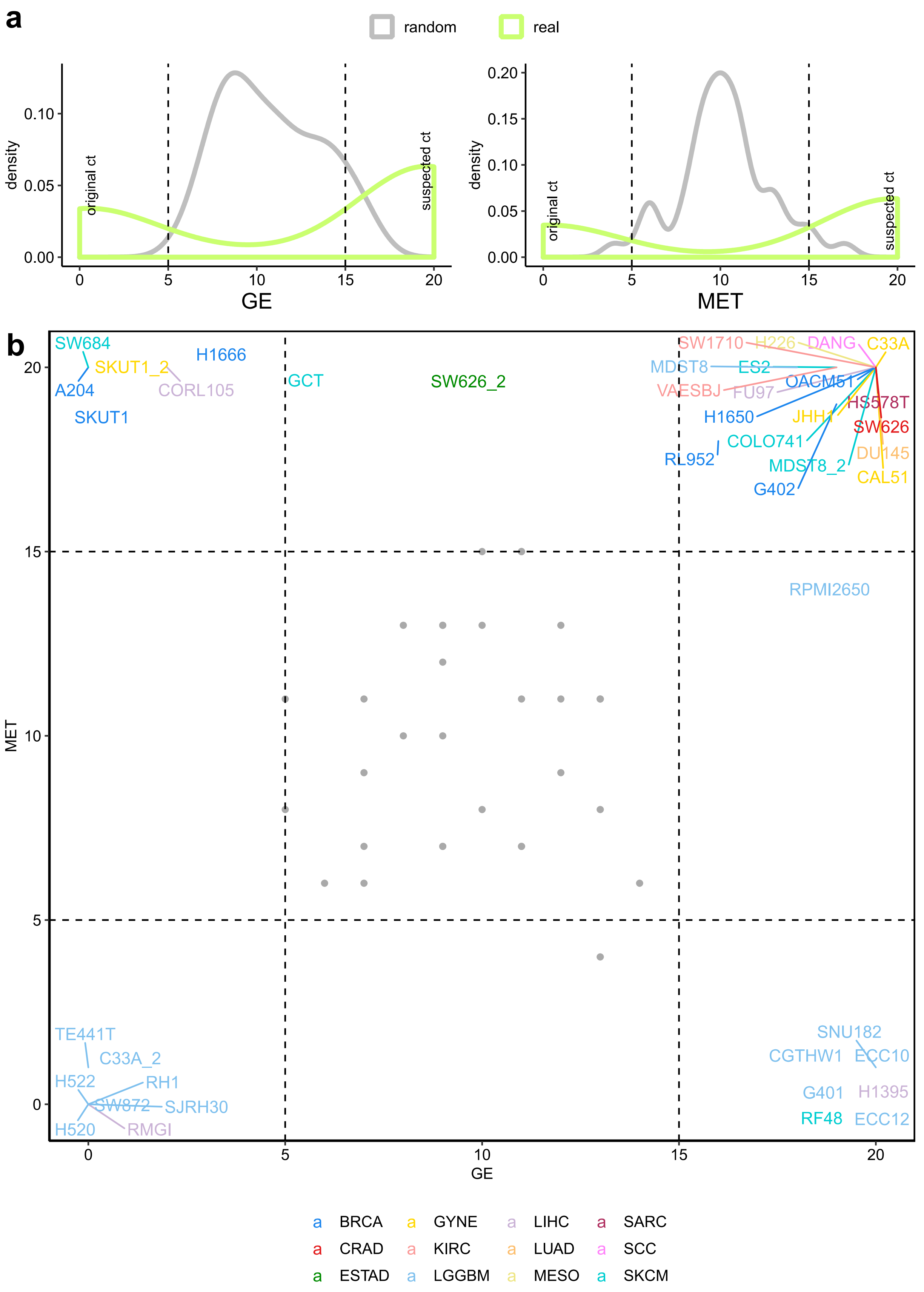


***Fig S5. (a)*** *Density of the prediction scores (classifier that predicts suspected versus original cancer type run 20 times) for GE and MET for the real model and a random model (same methodology but randomizing the labels before). A value of 20 means that it is predicted as suspected the 20 runs and a value of 0 means it is predicted as original the 20 runs.* ***(b)*** *Prediction scores for GE and MET for the suspected cell lines. Colors represent the suspected cancer type. Grey dots represent the random values.*

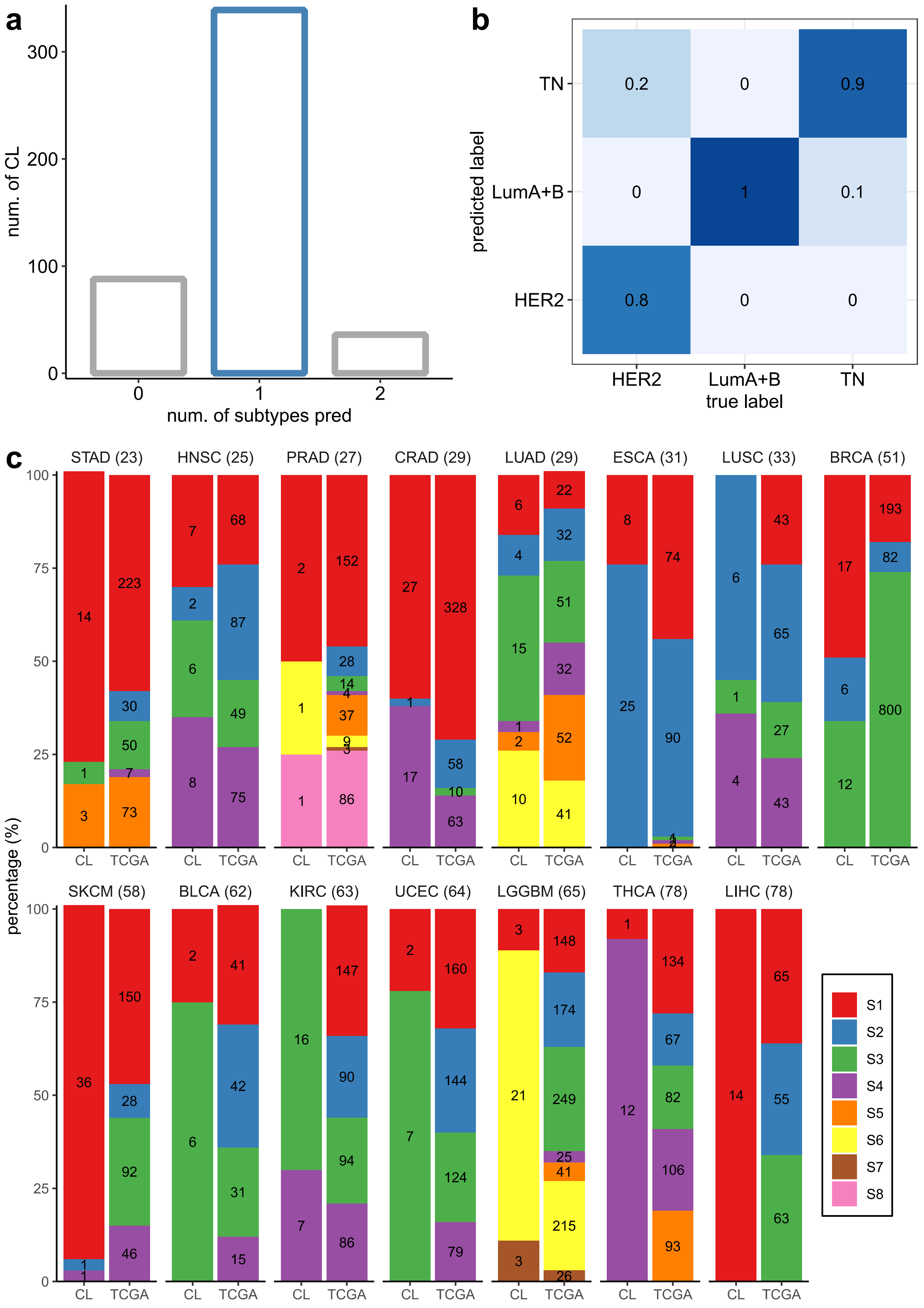


*Fig S6.* ***(a)*** *Number of cell lines that are assigned to 0, 1, 2 or 3 subtypes (used one-vs-rest subtype classifiers within each cancer type).* ***(b)*** *Normalized confusion matrix for the prediction of the breast cancer subtypes.* ***(c)*** *Subtypes proportions (raw counts shown in the bars) present in the cell lines (CL) and the human tumors (TCGA) for each cancer type. The cancer types are ordered from more to less similar according to euclidean distance between CL and TCGA percentages (shown in the title). Chi-square, p-value and euclidean distance for the differences between CL and TCGA in Supplementary Table 4.*

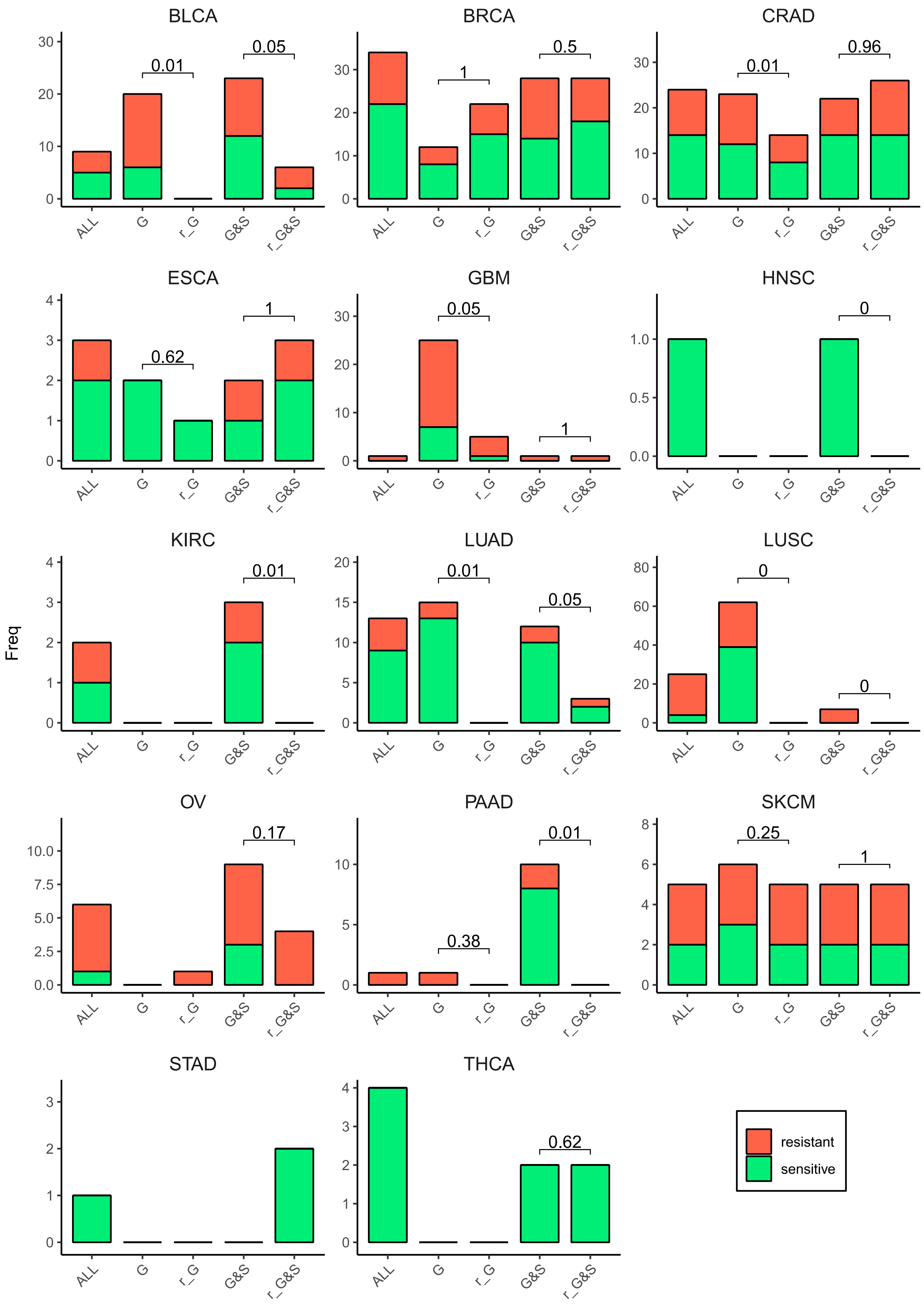


***Fig S7.*** *Number of significant associations between Cancer Functional Events (CFEs) and drugs detected in an ANOVA test for all cell lines (ALL), cell lines in the golden set (G), cell lines in the golden and silver set (G&S) and a random subset of cell lines that match the number of cell lines in the golden set (r_G) and in the golden and silver set (r_G&S). For the random subsets, the number of significant associations is calculated 10 times (randomizing the selection each time) and the number shown is the median of the 10 runs. P-value for a sign test (alternative = "less") between the associations in the G/G&S and the associations in the 10 runs of random_G/random_G&S are shown over the r_G/r_G&S bars.*

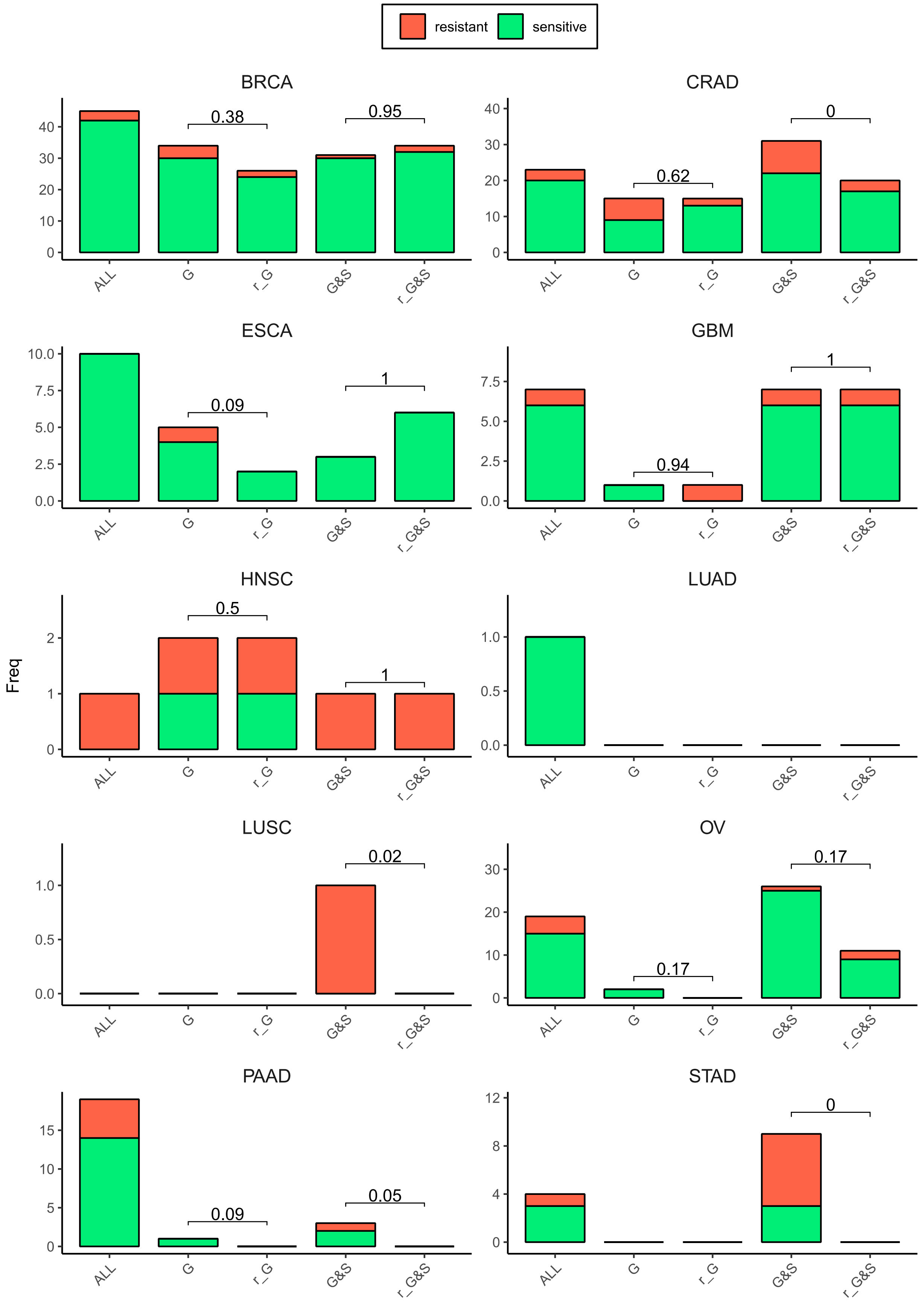


***Fig S8.*** *Number of significant associations between Cancer Functional Events (CFEs) and gene dependency (CRISPR k.o. genes) detected in an ANOVA test for all cell lines (ALL), cell lines in the golden set (G), cell lines in the golden and silver set (G&S) and a random subset of cell lines that match the number of cell lines in the golden set (r_G) and in the golden and silver set (r_G&S). For the random subsets, the number of significant associations is calculated 10 times (randomizing the selection each time) and the median of the 10 runs is shown. P-value for a sign test (alternative = "less") between the associations in the G/G&S and the associations in the 10 runs of random_G/random_G&S are shown over the r_G/r_G&S bars.*

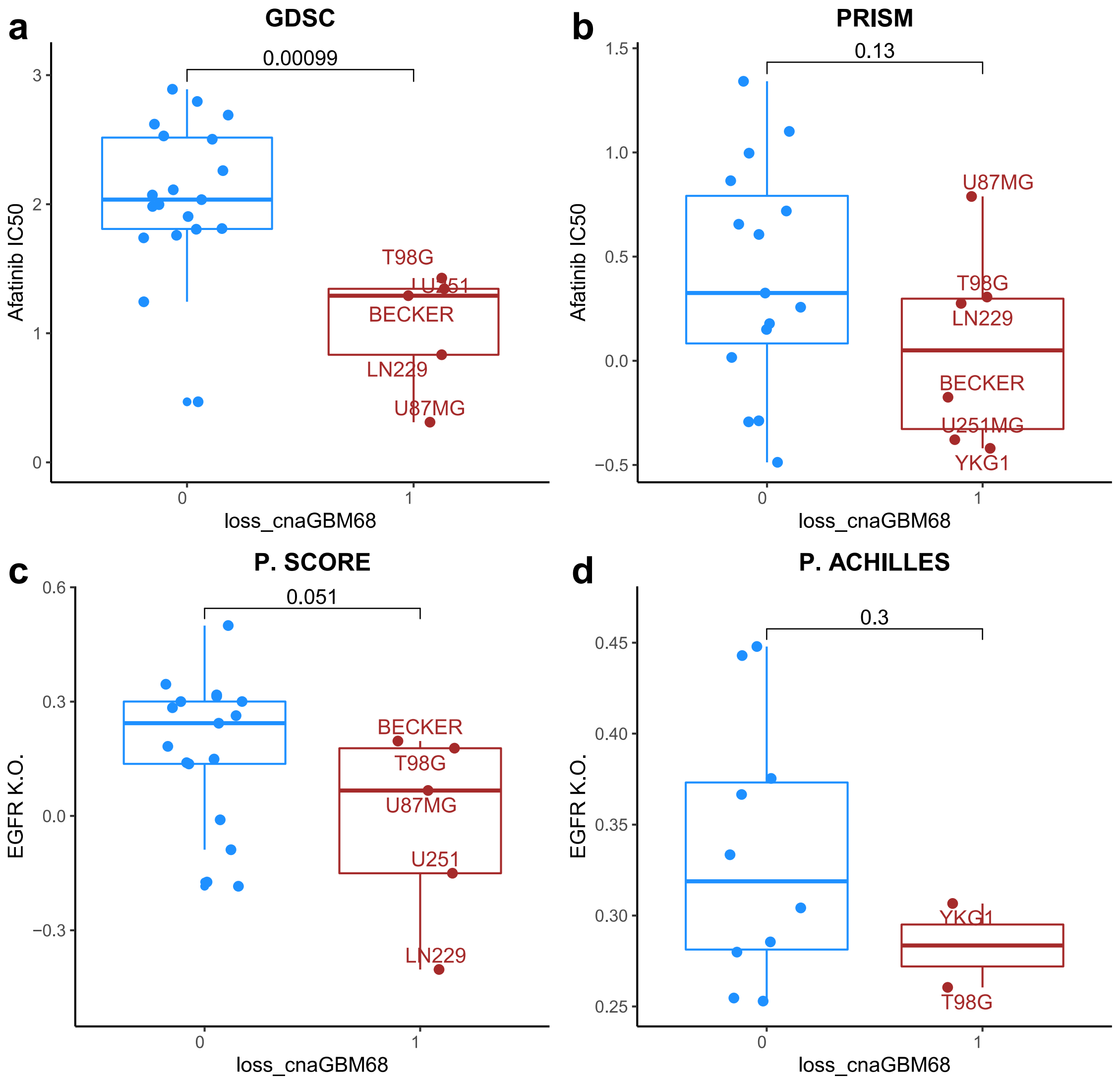


***Fig S9***. *Drug sensitivity (IC50) to afatinib in GBM cell lines (golden set only). Two groups are compared: cell lines with loss_cnaGBM68 (1) and without (0). IC50 obtained from GDSC* ***(a)*** *and from PRISM* ***(b).*** *Gene Dependency to EGFR K.O. in GBM cell lines (golden set only). Two groups are compared: cell lines with loss_cnaGBM68 (1) and without (0). Gene dependency data obtained from Project Score* ***(c)*** *and from Project Achilles (process with Project Score pipeline)* ***(d).*** *P-values calculated with wilcoxon-test and the alternative hypothesis “less”.*
