## Supplementary Figure S3 for "Matching cell lines with cancer type and subtype of origin via mutational, epigenomic and transcriptomic patterns"

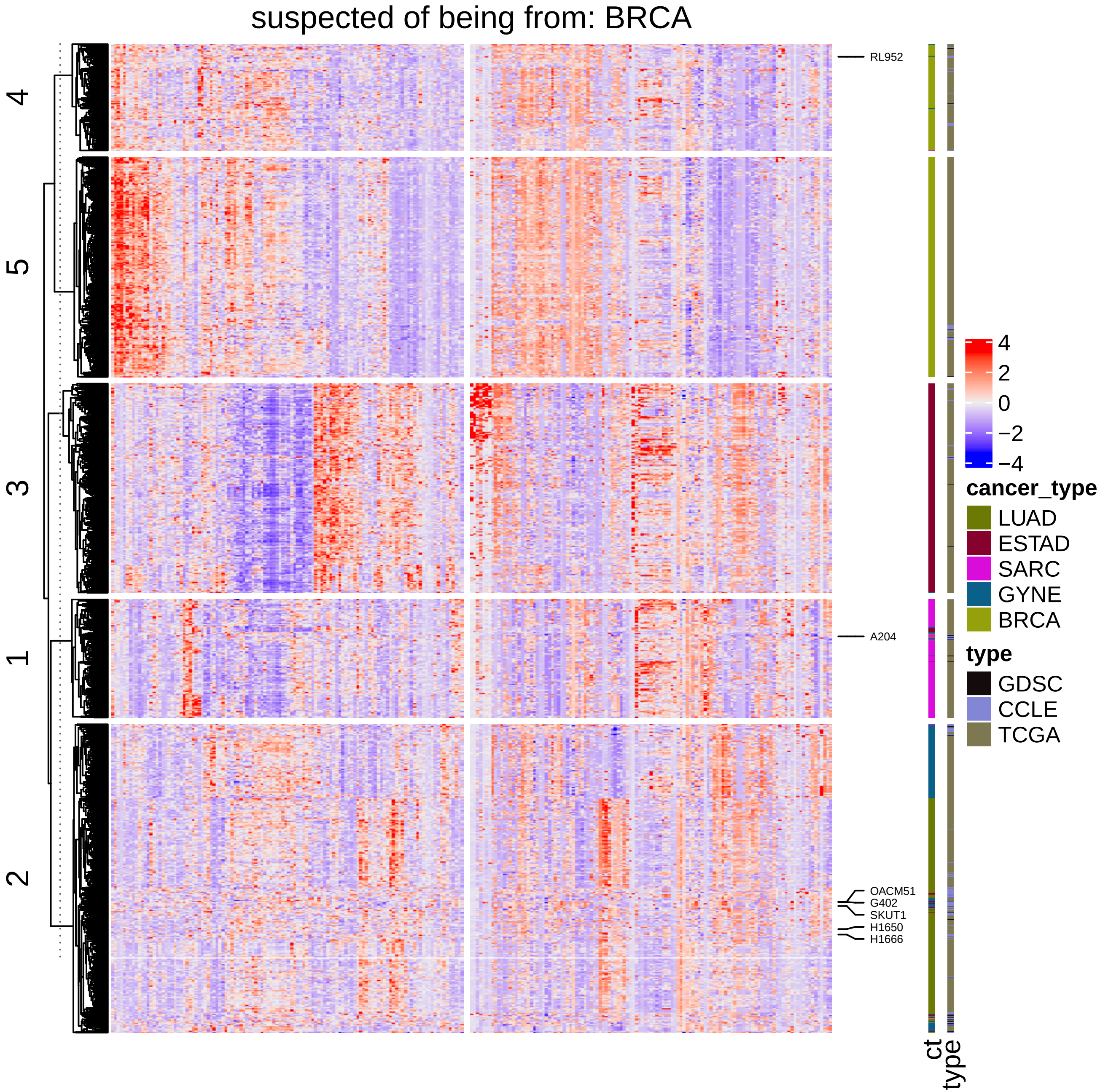

suspected of being from: BRCA
4
5
3
1
2
RL952
A204
OACM51
G402
SKUT1
H1650
H1666
4
2
0
−2
−4
cancer_type
LUAD
ESTAD
SARC
GYNE
BRCA
type
GDSC
CCLE
TCGA
ct
type

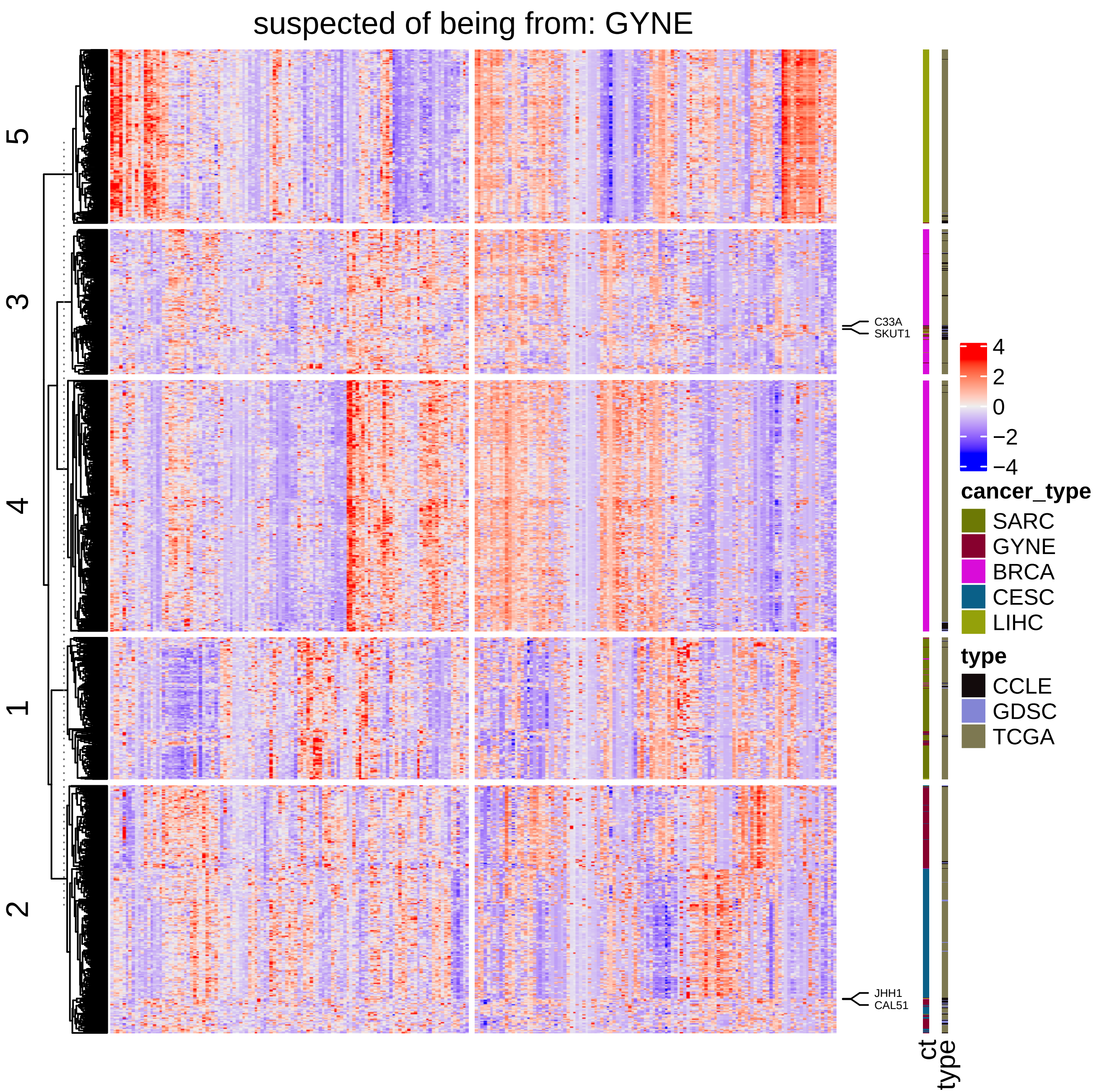
suspected of being from: GYNE
5
3
4
1
2
C33A
SKUT1
JHH1
CAL51
4
2
0
−2
−4
cancer_type
SARC
GYNE
BRCA
CESC
LIHC
type
CCLE
GDSC
TCGA
ct
type

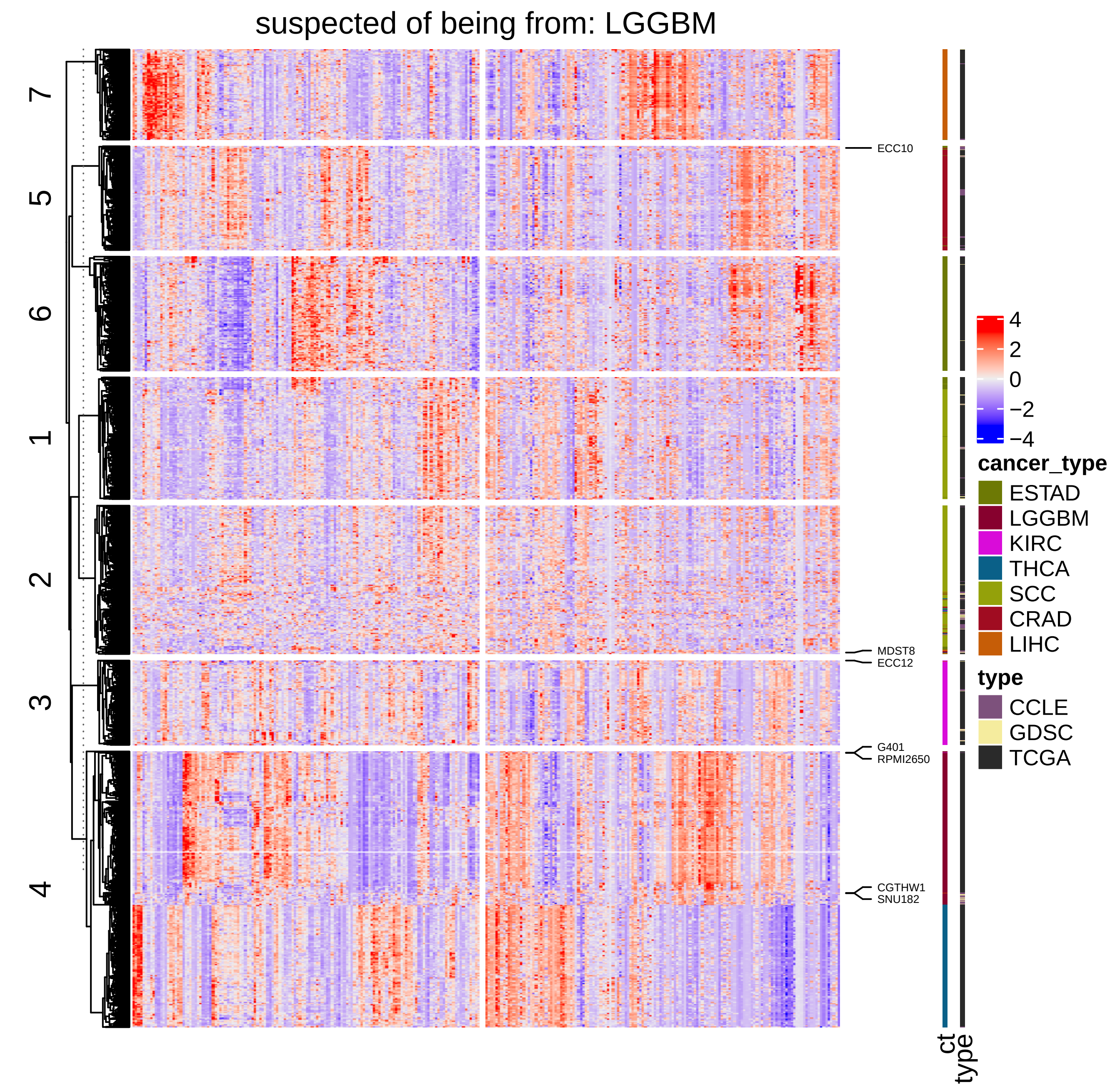

suspected of being from: LGGBM
7
5
6
1
2
3
4
ECC10
MDST8
ECC12
G401
RPMI2650
CGTHW1
SNU182
ct
type
4
2
0
−2
−4
cancer_type
ESTAD
LGGBM
KIRC
THCA
SCC
CRAD
LIHC
type
CCLE
GDSC
TCGA

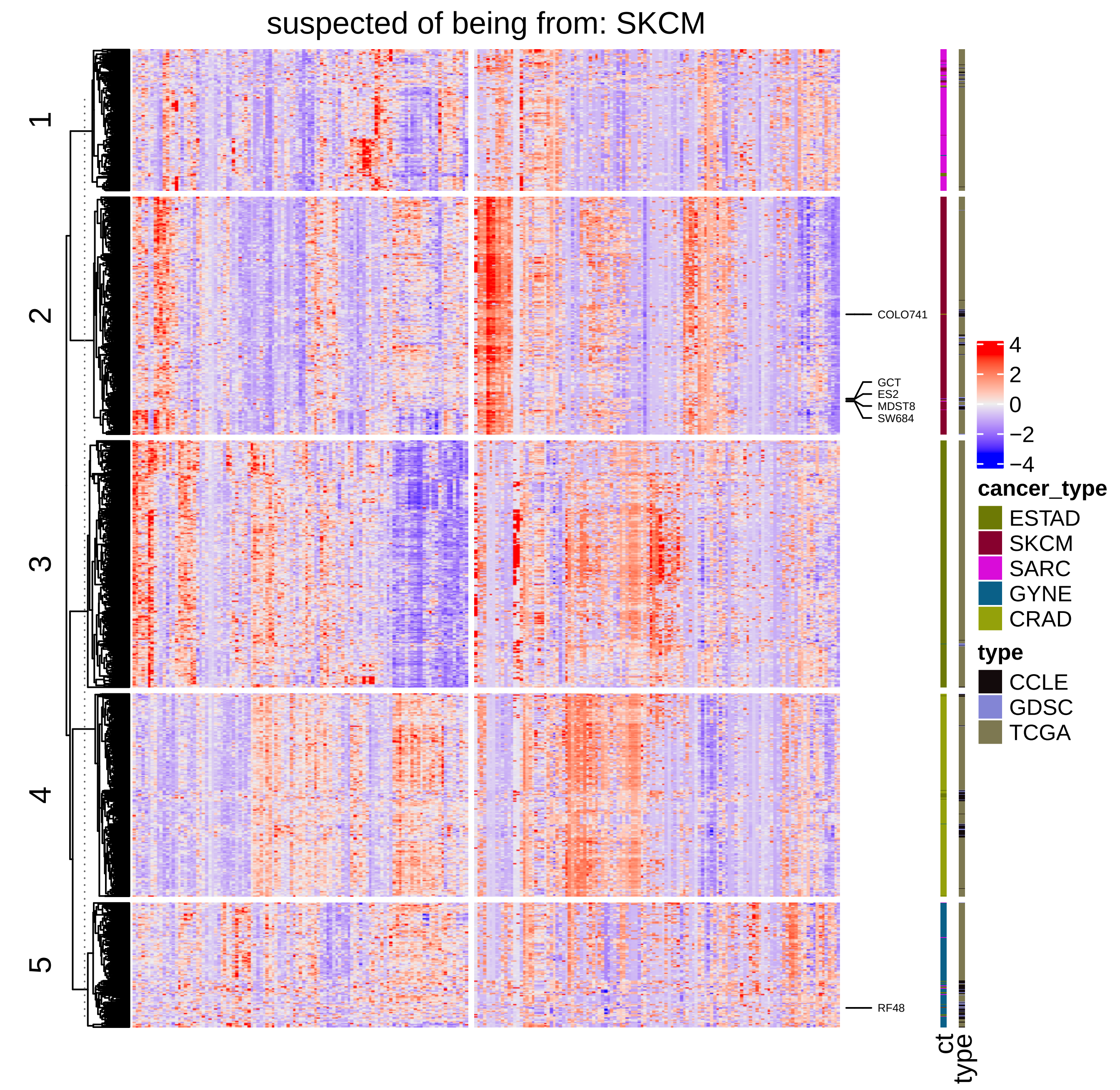
suspected of being from: SKCM
1
2
3
4
5
COLO741
GCT
ES2
MDST8
SW684
RF48
4
2
0
−2
−4
cancer_type
ESTAD
SKCM
SARC
GYNE
CRAD
type
CCLE
GDSC
TCGA
ct
type

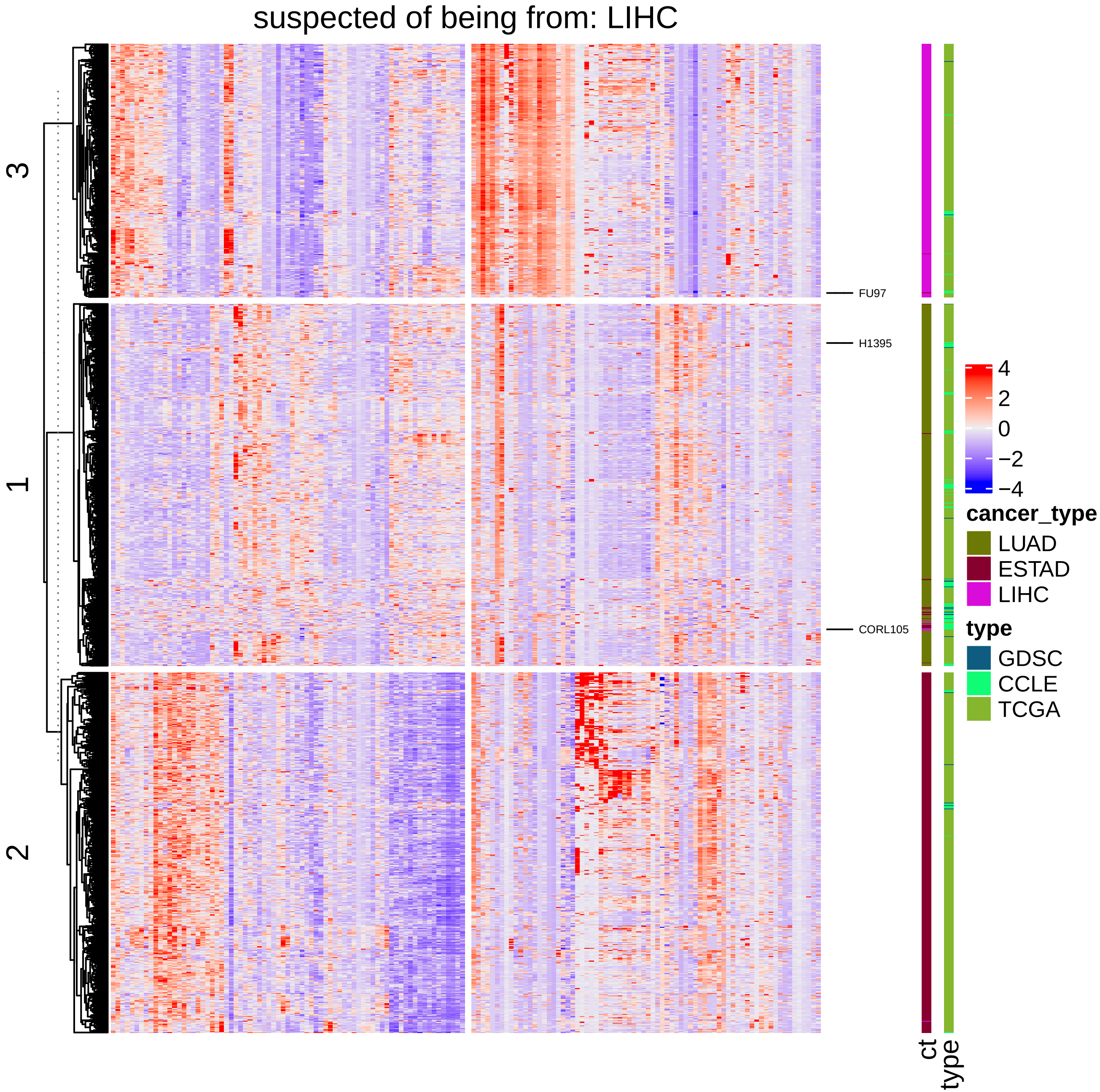

suspected of being from: LIHC
3
1
2
FU97
H1395
CORL105
ct
type
4
2
0
−2
−4
cancer_type
LUAD
ESTAD
LIHC
type
GDSC
CCLE
TCGA

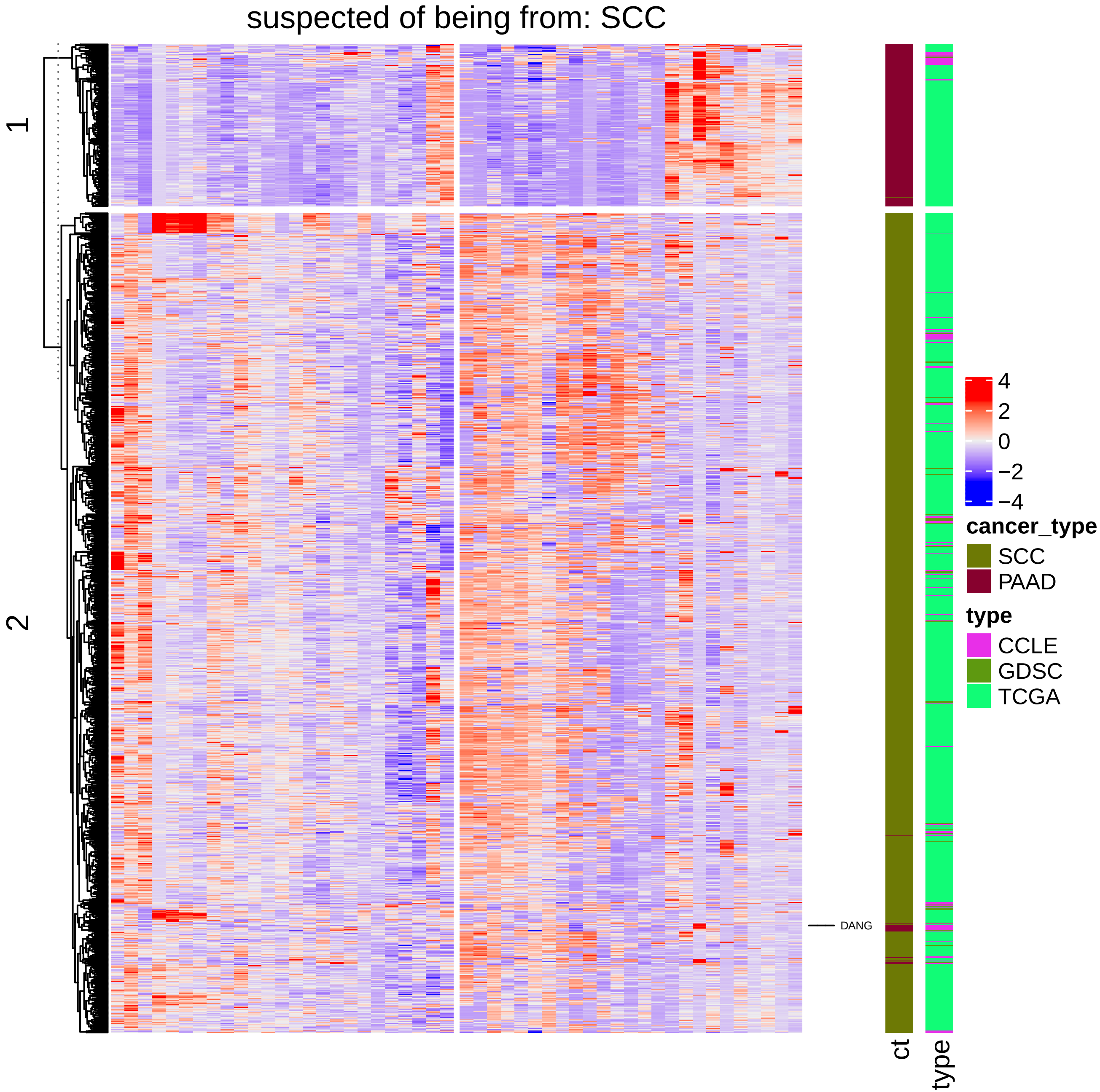

suspected of being from: SCC
1
2
DANG
4
2
0
−2
−4
cancer_type
SCC
PAAD
type
CCLE
GDSC
TCGA
ct
type

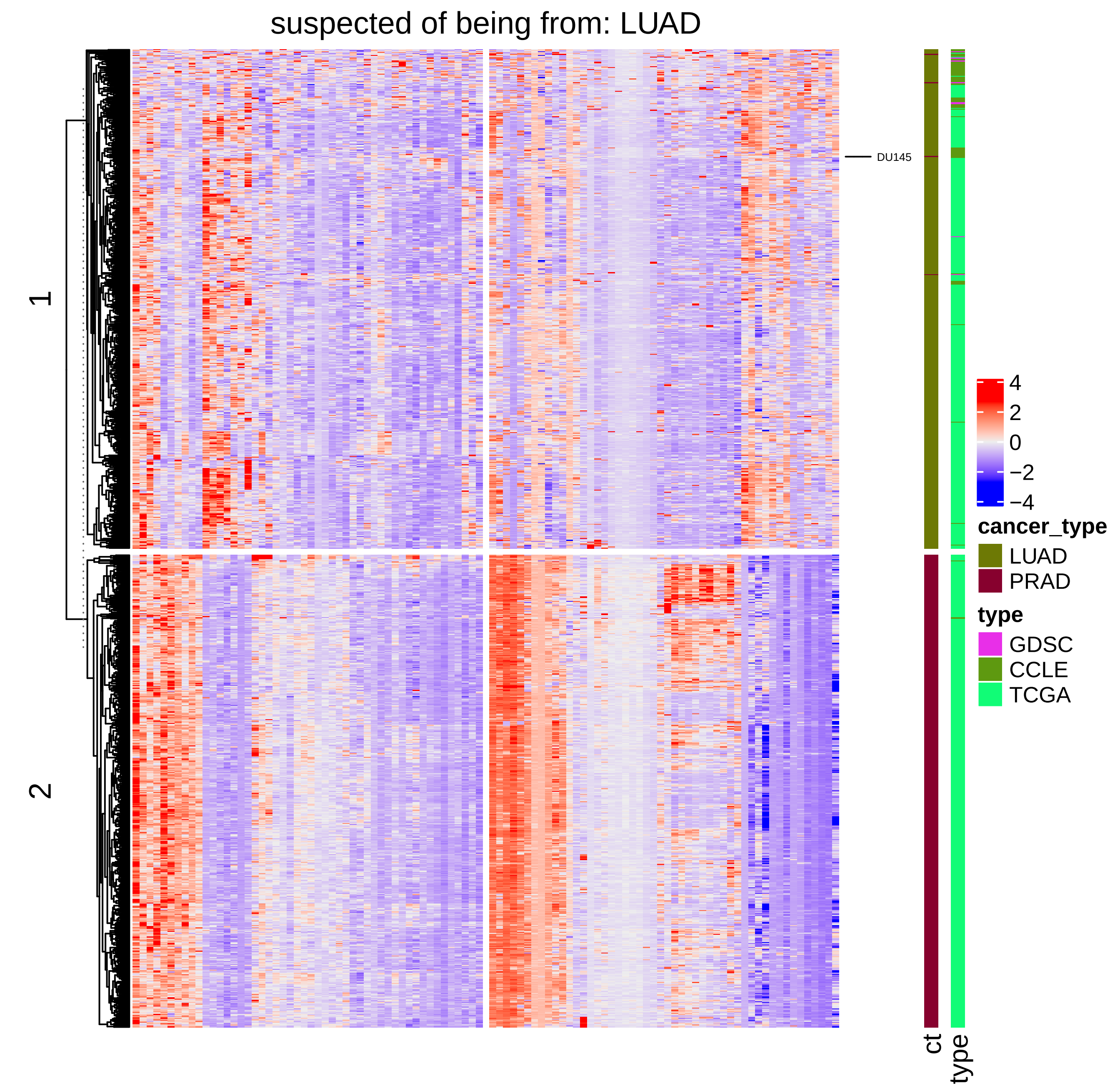
suspected of being from: LUAD
1
2
DU145
4
2
0
−2
−4
cancer_type
LUAD
PRAD
type
GDSC
CCLE
TCGA
ct
type

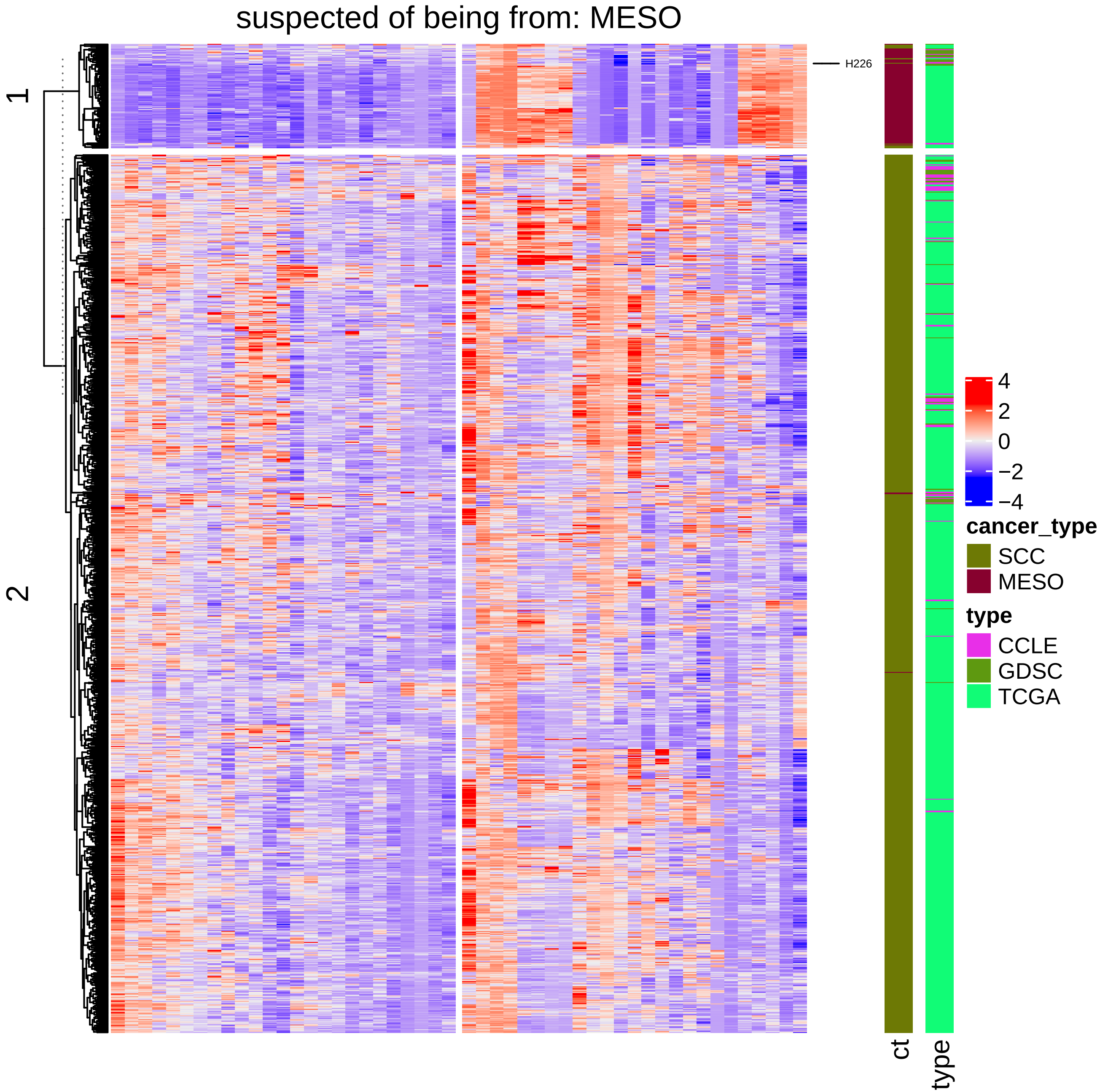

suspected of being from: MESO
1
2
H226
4
2
0
−2
−4
cancer_type
SCC
MESO
type
CCLE
GDSC
TCGA
ct
type

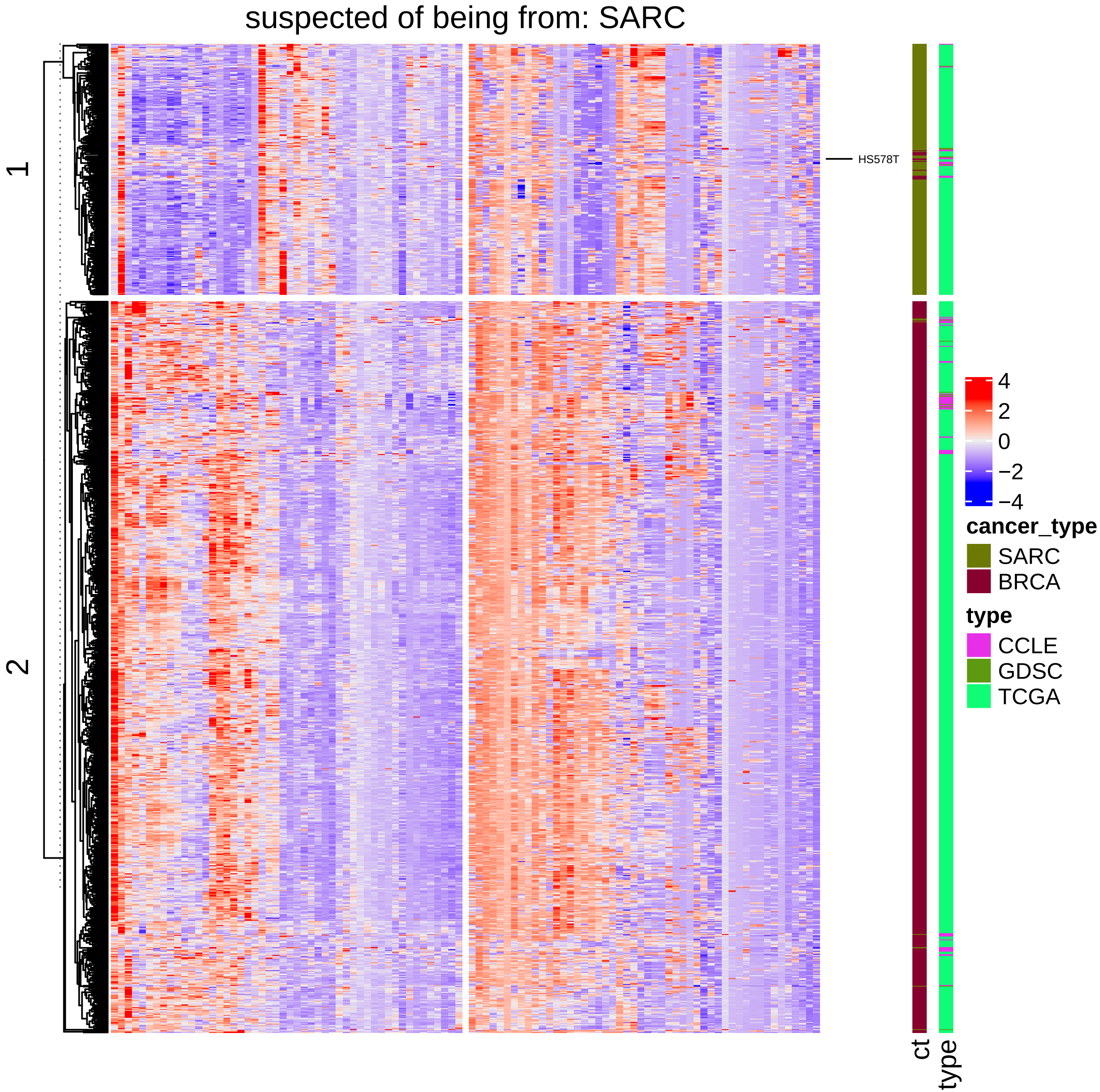
suspected of being from: SARC
1
2
HS578T
4
2
0
−2
−4
cancer_type
SARC
BRCA
type
CCLE
GDSC
TCGA
ct
type

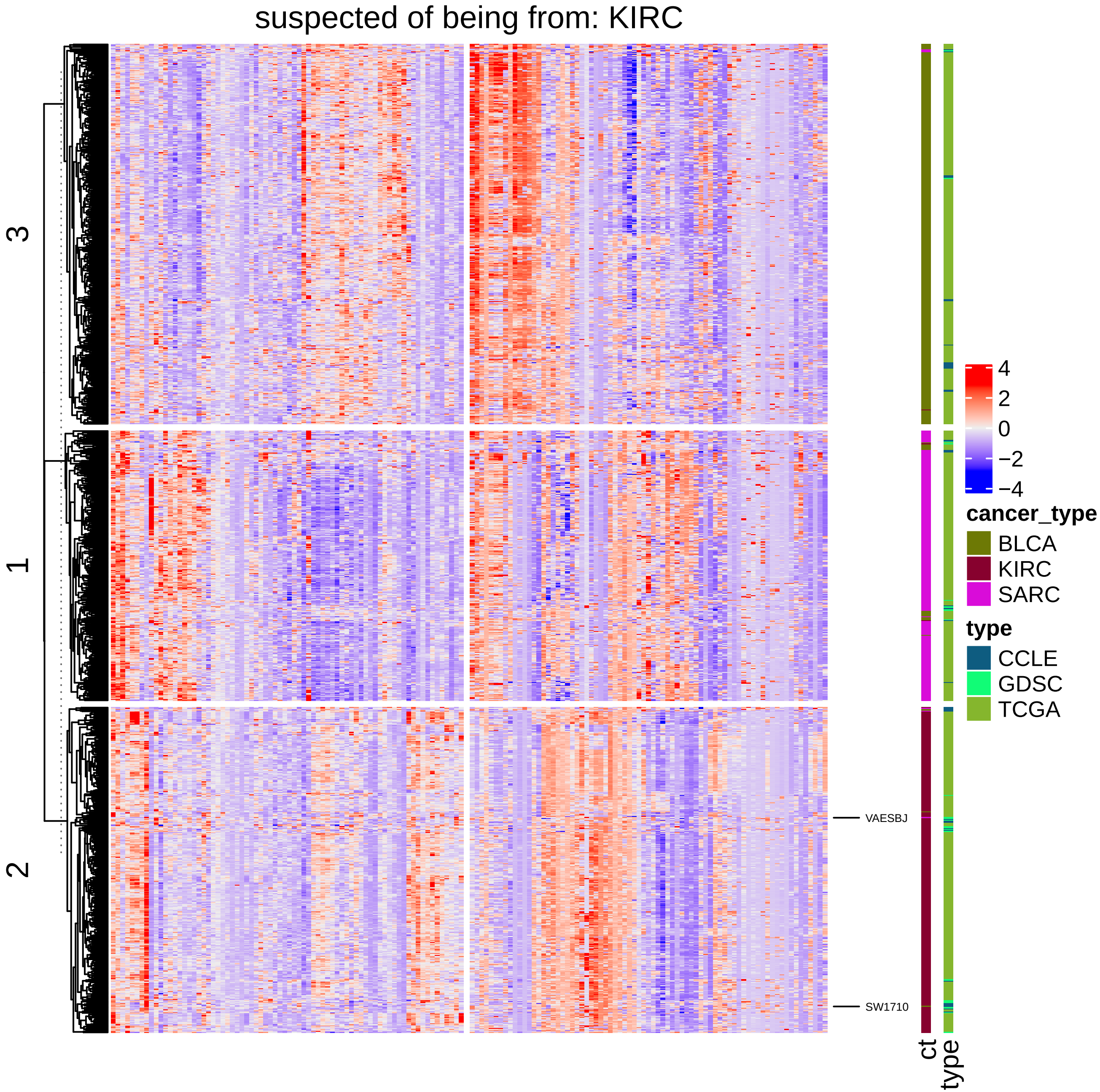
suspected of being from: KIRC
3
1
2
VAESBJ
SW1710
ct
type
4
2
0
−2
−4
cancer_type
BLCA
KIRC
SARC
type
CCLE
GDSC
TCGA

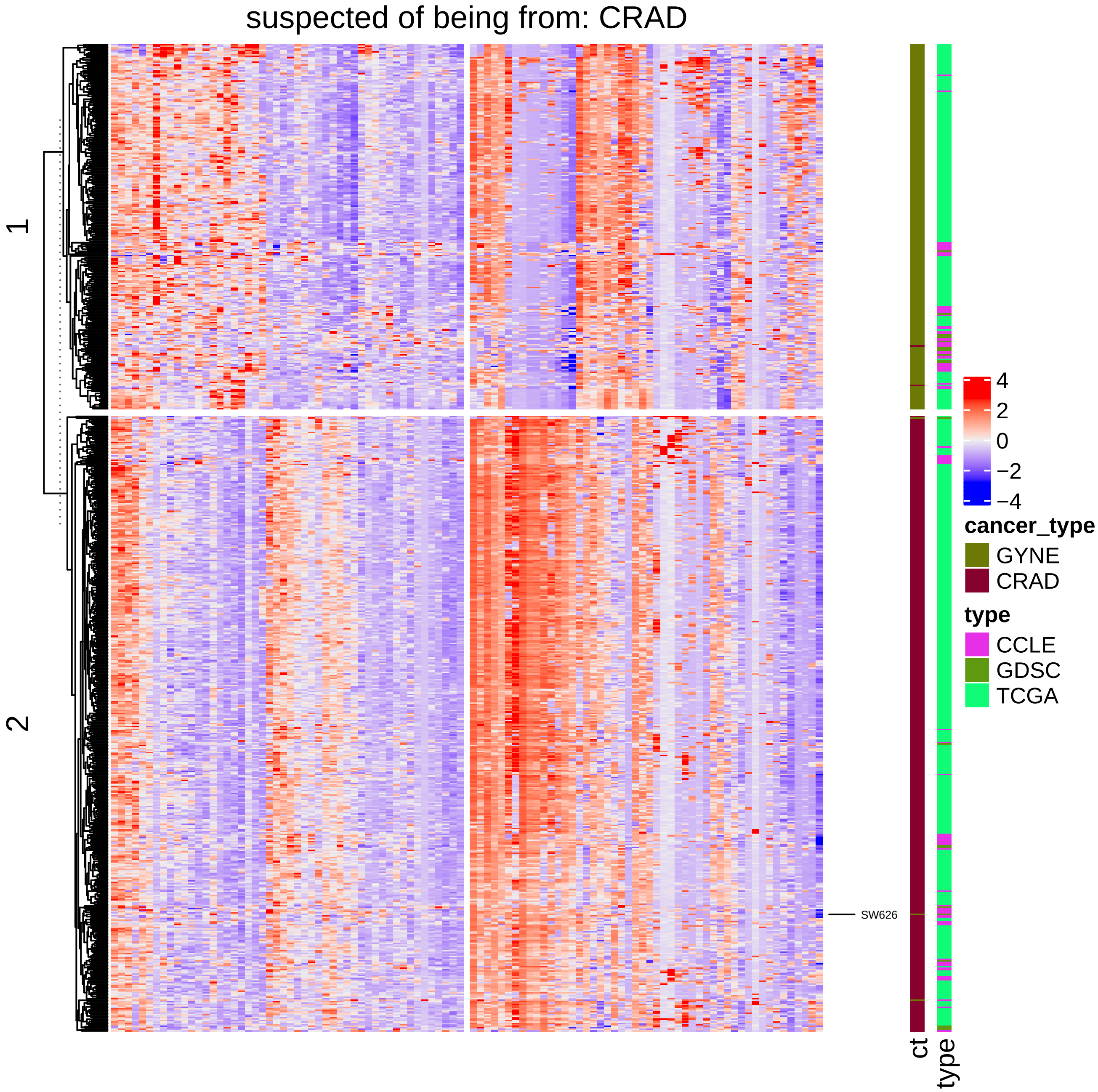
suspected of being from: CRAD
1
2
SW626
ct
type
4
2
0
−2
−4
cancer_type
GYNE
CRAD
type
CCLE
GDSC
TCGA

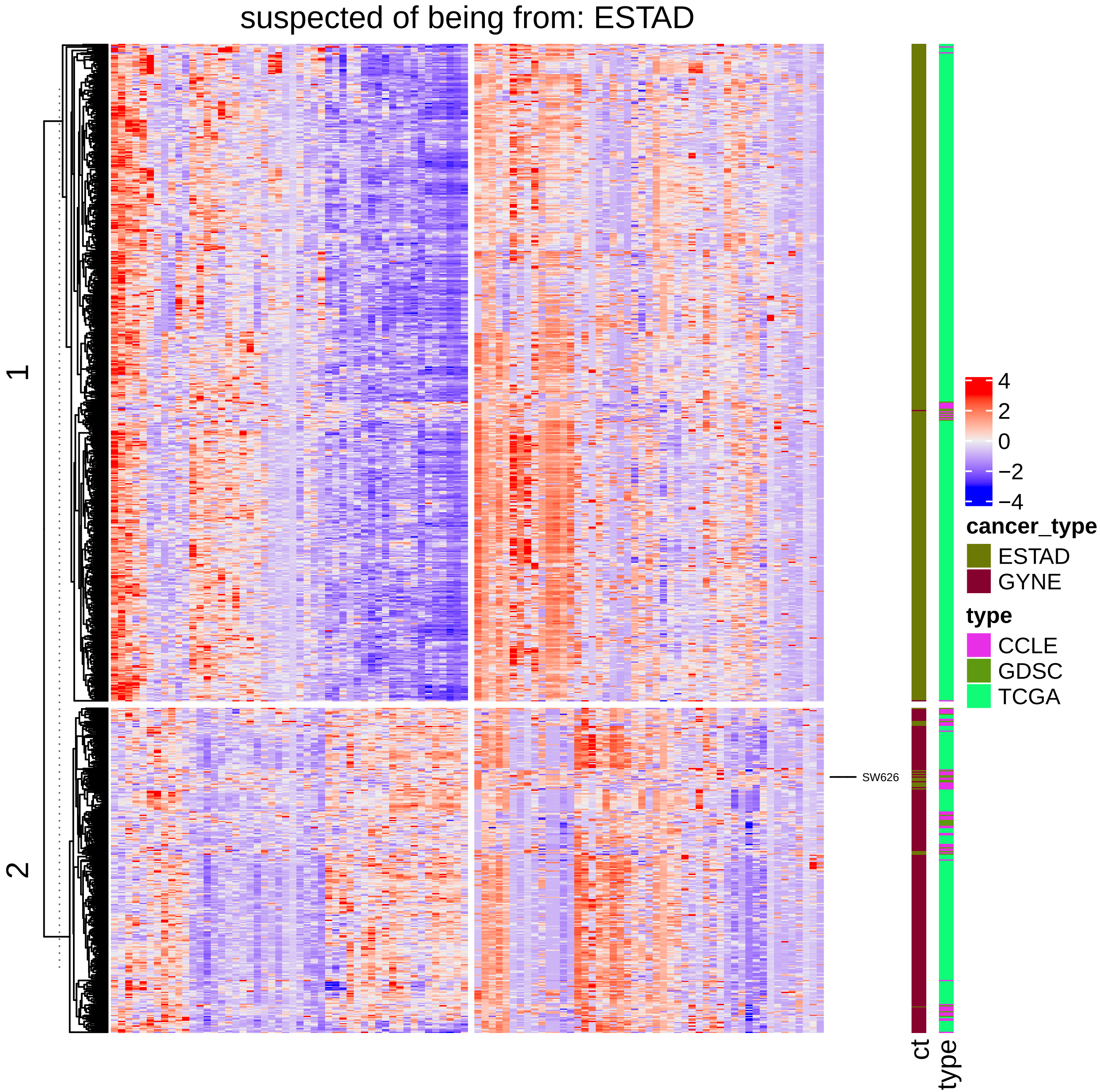

suspected of being from: ESTAD
1
2
SW626
ct
type
4
2
0
−2
−4
cancer_type
ESTAD
GYNE
type
CCLE
GDSC
TCGA
